# Electrochemically controlled blinking of fluorophores to enable quantitative stochastic optical reconstruction microscopy (STORM) imaging

**DOI:** 10.1101/2023.09.13.557504

**Authors:** Ying Yang, Yuanqing Ma, Richard Tilley, Katharina Gaus, J. Justin Gooding

## Abstract

Stochastic optical reconstruction microscopy (STORM) allows widefield imaging with single molecule resolution through calculating the coordinates of individual fluorophores from the separation of the fluorophore emission in both time and space. Such separation is achieved by photoswitching the fluorophores between a long lived OFF state and an emissive ON state. Despite STORM having revolutionized cellular imaging it remains challenging for quantitative imaging of single molecules due to a number of limitations, such as photobleaching caused under counting, overlapping emitters related fitting error, and repetitive but random blinking induced over counting. To overcome these limitations, we develop an electrochemical approach to switch the fluorophores between ON and OFF states for STORM (EC-STORM). The approach provides greater control over the fluorophore recovery yield, emitter density, and random blinking than photochemically switching. The result is EC-STORM has superior imaging capability than conventional photochemical STORM and can perform molecular counting; a significant advance.

## Main Text

Super-resolution light microscopy methods have revolutionized cell biology by enabling imaging with resolution below the diffraction limit of light ^1-3^ such that of single proteins and cellular structures can be observed in live cells^4-9^. The suite of SMLM methods, including stochastic STORM, photo-activated localization microscopy (PALM) and point accumulation for imaging in nanoscale topography (PAINT) ^10-15^ are particularly attractive because they use a conventional widefield approaches that provide one order of magnitude higher resolution^16^. This is achieved via the minimization of the overlap of the point spread function of the emission from individual fluorophore in a given image frame. A super-resolution image is generated by reconstructing the fluorophore locations over many imaging frames^15, 17^. To achieve this requires a small subset of fluorophores to be in their fluorescent ON state at each given frame. In the STORM technique an organic dye is photochemically switched into an OFF state using a high-power visible laser. Either spontaneously, or photoinduced with a second UV laser, a sparse subset of dyes is then switched to the ON state and their positions can thus be precisely determined. Since a STORM image is built molecule by molecule, it should be possible to determine the numbers of underlying molecules. However, achieving this has proven difficult due to the limitations existing in imaging and localization steps, as well as the photophysical behavior of the dyes. For example, the initial off switching of the dyes with an intense laser leads to photobleaching, the bleached molecules will no longer be visible and result in undercounting. The overlapping of fluorophores often occurs with densely labelled biological samples which can lead to missed events and undercounting in diffraction limited volume. While the most problematic is the multiple blinking problem of organic dyes in STORM, making a single dye appear as several molecules. This issue is especially difficult to overcome for some cyanine-based dyes, like Alexa 647, for which the dark time between re-blinking can be too long for accurate temporal grouping.^18^

We address these challenges using electrochemical switching of the fluorophore. The inspiration for exploring electrochemical switching came from the previous research that the fluorescence of some fluorophores can be altered by external electrical potential.^19-21^ Our recent study showed that the fluorescence intensity of Alexa Fluor 647 (Alexa 647) could be modulated by the electrochemical potential (Supplementary Figure 1-3, Supplementary Movie 1)^22^. However, in previous research^19-22^ only the brightness of the fluorophore can be tuned using electrochemical potential, the stochastic switching of the molecules between their ON and OFF states as required for STORM imaging was not achieved. To understand why electrochemical switching of fluorophores might be possible requires an understanding of how the fluorophores can be photochemically switched between the ON and OFF states. A simplified Jablonski diagram (Fig. 1a) for the commonly used STORM dye Alexa 647 depicts the mechanism^10,23,24^. Briefly, the OFF state transfer of Alexa 647 is a photoinitiated thiol–ene reaction^15,25^. The majority of the Alexa 647 dyes are firstly excited into a singlet state using a 642 nm laser at high power (with laser intensity of 10-30 kW cm^-2^) where they can further intersystem cross into a triplet state ^26-28^. From the triplet state the fluorophores can then enter a long-lived dark state^15^ when organothiols reacts with the polymethine bridge of Alexa 647 and disrupts the conjugated π-electron system^25,29,30^. The reaction is reversible such that the C-S bond for the OFF-state Alexa 647 can dissociate, either by oxidation by oxygen or by UV irradiation, which will switch Alexa 647 back to the ON state. In conventional STORM, to keep Alexa 647 in their OFF states longer, oxygen scavenging system like glucose and glucose oxidase is widely used to reduce the oxygen concentration by ten times with ∼5-20 μM.^31^ During STORM image acquisition, the oxygen and thiol concentrations in the buffer solution, together with the intensities of the imaging and activation lasers, can be further altered to optimize the number of fluorophores in the ON state at any given time.

**Fig. 1.**
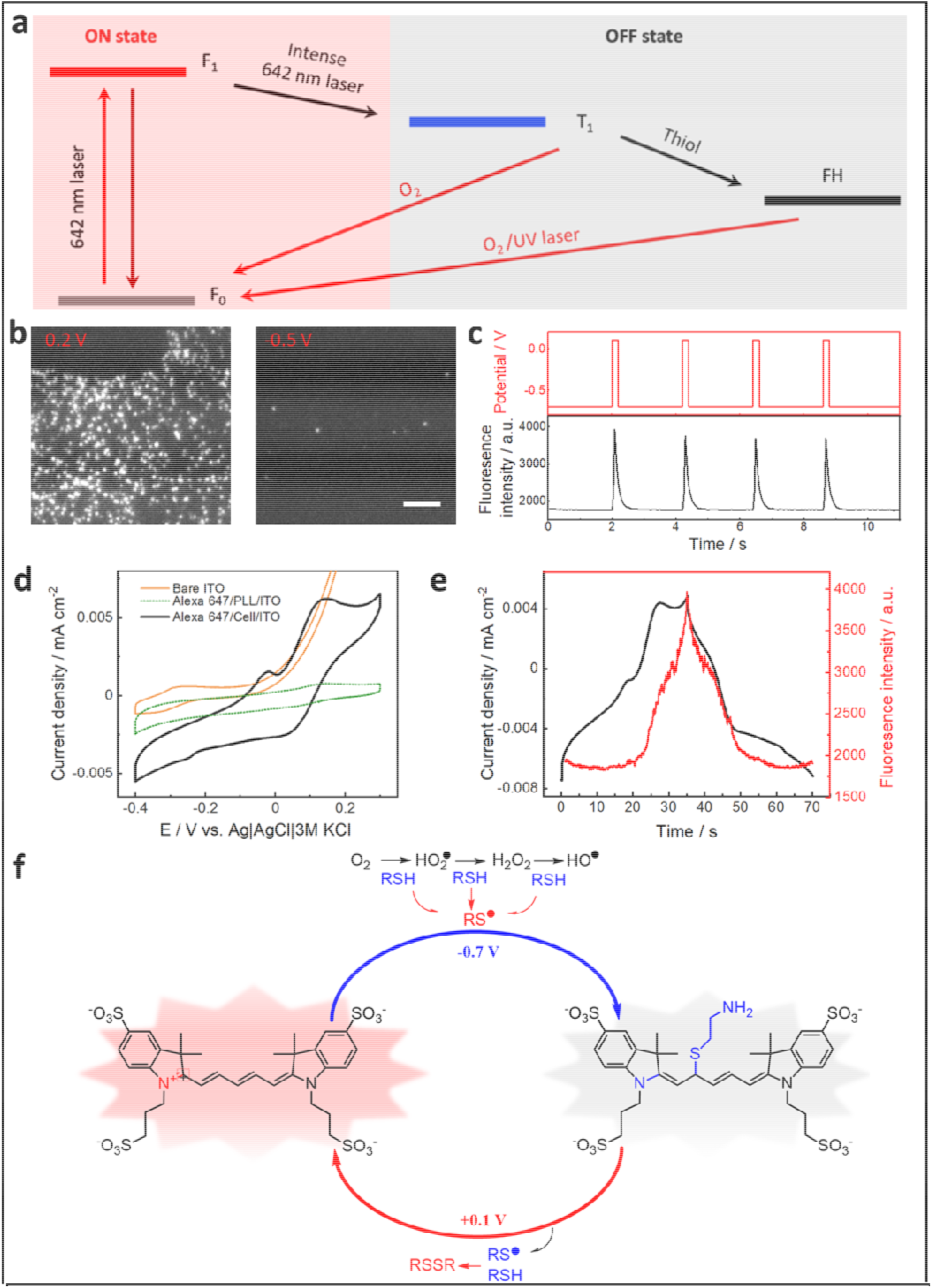
Electrochemical switching fluorescence of Alexa 647. **a**, Simplified Jablonski diagram showing how Alexa 647 is switched between the ON and OFF state under STORM conditions. In the bright ON state, Alexa 647 can be excited from the ground state (F_0_) to the excited singlet state (F_1_). From F_1_, Alexa 647 may either relax to the ground state by emitting photons or alternatively undergo intersystem crossing to the dark triplet state (T_1_). From the triplet state, molecules may return to the ground state via oxidation by oxygen or progress to a long-lived dark state (FH) after reacting with a primary thiol. Alexa 647 molecules in the dark state may return to the ground state by oxidation by oxygen, or alternatively, by exposure to near-UV radiation. F_0_ and F_1_ are called ON states, while T_1_ and FH are OFF states. At ambient oxygen levels, T_1_ can last microseconds, whereas FH can last between milliseconds and minutes. **b**, Images frame of Alexa 647 labelled microtubules (α-tubulin) of COS-7 cell in oxygen scavenger tris buffer with 50 mM cysteamine, a common STORM buffer, that shows a subset of Alexa 647 stochastically blinking. The frames show that at a potential of 0.2 V the majority of Alexa 647 are in their ON state whilst at -0.5 V many of the Alexa 647 molecules appear to be in OFF state. A 642 nm laser with intensity of 2 kW cm^-2^ at the back aperture of the objective was used as the excitation source during imaging. **c**, the corresponding electrochemical potential regulated fluorescence switching curve for b. The imaging sample was COS-7 cells, where the microtubules were pre-labeled with primary anti α-tubulin antibodies and then with secondary anti-rabbit IgG antibodies tagged with Alexa 647. The scale bar is 5 μm. **d**, Cyclic voltammograms of bare ITO (blue dashed line), Alexa 647/PLL/ITO (green dotted line), and Alexa 647/Cell/ITO (black line) in the STORM buffer at a scan rate of 20 □ mV □ s^−1^. The Alexa 647/PLL/ITO sample is prepared by directly absorbing Alexa 647 on poly-L-lysine coated ITO, the Alexa 647/Cell/ITO sample is COS-7 cells with Alexa 647 labeled microtubules attached ITO surface. **e**, the correlation between the fluorescence intensity and current change corresponding to the redox peaks at 0.17 V/0.08 V on cyclic voltammograms for Alexa 647/Cell/ITO sample. **f**, Proposed mechanistic pathways for electrochemical induced switching of Alexa 647 between its ON and OFF states. Under negative potential, the thiol reacts with the -ene functional group at Alexa 647 via an anti-Markovnikov addition to form a thiol ether, and hence the Alexa 647 will be switched into their OFF state. Under positive potential, the C-S bond dissociation will switch the Alexa 647 back to the ON state.

To assess whether we could use electrochemistry to control the thiol-ene reaction between thiol and Alexa 647. We used an indium tin oxide (ITO) coated glass coverslip as the microscope slides to perform electrochemistry and fluorescence imaging simultaneously^22,32-34^ (Supplementary Figure 1). The standard oxygen scavenger tris buffer with cysteamine,as conventionally used for STORM imaging with Alexa 647, was utilized as imaging and electrochemical buffer solution. As can be seen from Fig. 1b-c and Supplementary Figure 5, the Alexa 647 can be switched from its ON and OFF by applying positive and negative electrochemical potentials on ITO surface in the STORM buffer respectively. The fluorescence intensity observed relates to the number of Alexa 647 molecules in the ON state. Much more ON state Alexa 647 can be observed at a positive potential (+0.2 V) than a negative potential (−0.5 V). Supplementary Movie 2 clearly shows that at constant potential, Alexa 647 molecules are exhibiting the characteristic stochastic blinking behavior, while the external electrochemical potential has a direct effect on the emitter density. That is, only the 642 nm excitation laser at moderate intensity (2 kW cm^-2^) was required, and the emitter density can be tuned separately using the electrochemical potential now.

To understand the underlying electrochemical reactions between the OFF and ON state transfer of Alexa 647, electrochemical characterizations using cyclic voltammetry (Fig. 1d, e) and different pulsed voltammetry (Supplementary Figure 6a) was performed. We note that the redox peaks at 0.17 V/0.08 V on cyclic voltammograms are most likely the electrochemical thiol-ene reaction between thiol and Alexa 647. Evidence for this comes from the fact that these peaks only appeared when both thiol and Alexa 647 were presented. More importantly, the fluorescence change correlates nicely with the current change in the potential range between 0 V and 0.3 V with representative redox peaks at 0.17 V/0.08 V (Fig. 1e and Supplementary Movie 3) The proposed overall reaction, and a possible radical involved mechanism, is given in Fig. 1f. When the potential is scanned to negative direction, several oxygen reduction reactions may occur, with reactive oxygen species being formed in aqueous solution^35-37^. The intermediate oxygen radicals or products, such as the extremely reactive hydroxyl radical (HO^•^)^38, 39^, hydroperoxyl radical (HO_2_ ^•^) and H_2_ O_2_ ^40^ can react with organothiols to generate thiyl radicals^41^. Thiyl radicals can react with ene-functional group at a negative potential in the thiol-ene click reaction. For stochastic switching of the fluorophores to occur however implies a very small amount of this reaction occurs in a stochastic manner. When a constant potential was applied on the ITO surface, we observed that the more negative the potential, the more Alexa 647 on the surface was converted to the OFF state. We proposed this is due to the more oxidative oxygen intermediates generated that promote more thiyl radical formations, followed the more Alexa 647 was switched to the OFF state by thiol addition. For the reverse reaction, we propose the C-S bond dissociation occurs during the electrochemical oxidation of the thiol adducted Alexa 647^42^. The dissociated thiol or its radicals will quickly react with other thiols to form a disulfide rather than propagating with the -ene functional groups at Alexa 647 as the oxidation reactions are thermodynamically more favourable under a positive potential.

Having ascertained that electrochemical potential can be used to switch the fluorophores into the OFF state and back to the ON state we next sought to understand the level of control electrochemistry gave us over the imaging parameters, considering the precise control we have over the potential that can be applied to the ITO. Fig. 2a and Supplementary Figure 7 show that the emitter density in the field of view increases with an increase in electrochemical potential. Alternatively, pulsing the potential to -0.7 V resulted in almost complete suppression of the number of Alexa 647 emitters in the ON state. The control over the emitter density is shown in Fig. 2b where emitter densities as low as 0.05 molecule per μm^2^ and as high as 1.2 molecule per μm^2^ are achieved. In contrast, with conventional STORM on the same sample, emitter densities only vary between 0.6 molecule per μm^2^ and 1.2 molecule per μm^2^.

**Fig. 2.**
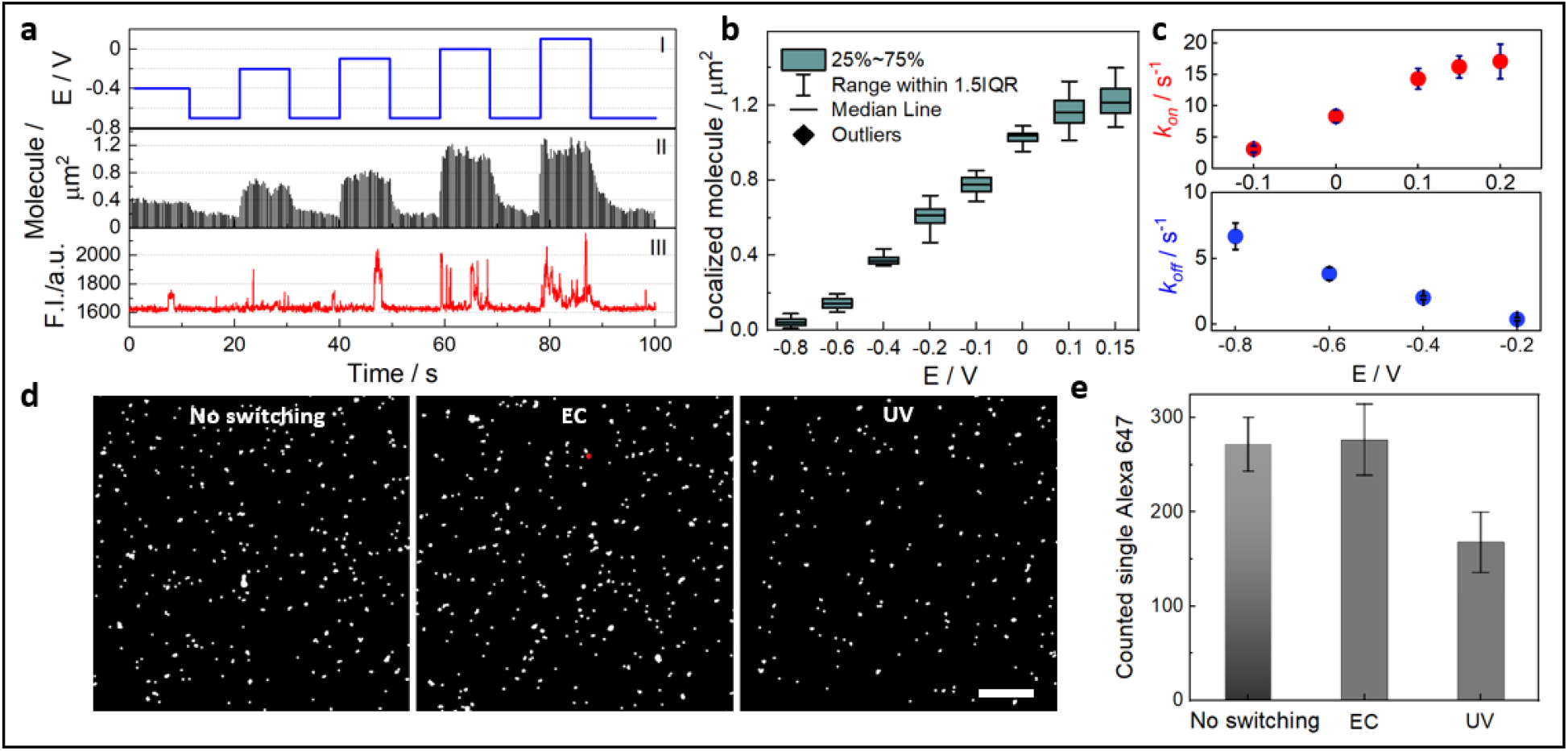
Study of emitter density, switching kinetics and recovery yield of Alexa 647. **a**, Time series of alternating pulses of electrochemical potentials showing reversible switching of the ON and OFF states of the Alexa 647 on microtubules in COS-7 cells, as determined from the number of localization events at each potential. (I) The applied potential versus time curve, on which the potentials are switched from -0.7 V to -0.4, -0.2, -0.1, 0, and +0.1 V respectively for 10 s at each potential. (II) The corresponding ON-state Alexa 647 density at each applied potential. The data was extracted from Supplementary Movie 4. (III) Fluorescence intensity trajectory in one pixel showing the higher probability of the Alexa 647 being in the ON state at more positive potentials, while most of time, the Alexa 647 in this region are in the OFF state at negative potentials. **b**, Box plot for the density of localization events at each applied potential. At positive potentials greater than 0.1 V, the density of localization events reached plateau as overlapping emitters could no longer be localized. At the open circuit potential (at around -0.2 V), the emitter density represents the same density as observed in conventional STORM condition. The data was extracted from Supplementary Movie 5. **c**, Fluorescence switching ON and OFF rates obtained by fitting localized molecule-time curves when switching electrochemical potentials between positive and negative values. **d**, Reconstructed STORM images obtained under different conditions with Alexa 647 molecules adsorbed directly onto a poly-L-lysine modified ITO slide. The presented images come from different regions of the same slide (see Method for sample preparation). A total of 2,000 image frames were collected to reconstruct each STORM image. ‘No switching’ refers to a STORM-style image performed in oxygen scavenger buffer but without cysteamine such that all the adsorbed Alexa 647 molecules are expected to remain in the ON state. The setting for ‘EC’ and UV’ conditions were detailed in Method. The scale bar is 5 μm. **e**, The corresponding recovered molecules that could be imaged under ‘no switching’, ‘EC’, and ‘UV’ conditions. Error bars in c and e denote mean ± standard deviation (n = 6 replicates of distinct samples).

Supplementary Movie 4 illustrates the dynamic switching when altering the applied potential between negative and positive values saw the number of Alexa 647 molecules blinking being a function of the applied potentials. The number of emitters is related to the rate constants for the switching of Alexa 647 between the fluorescence ON and OFF states (*k*_*on*_ and *k*_*off*_). We compare the rate constants from electrochemical switching with the rates determined using the conventional photoswitching pathway in typical STORM experiments (see Method). With conventional STORM (Supplementary Figure 8), *k*_*on*_ increases with the intensity of UV laser and *k*_*off*_ increases with the intensity of the 642 nm laser, as others have found^15, 43^. Fig. 2c shows that with electrochemical switching that *k*_*on*_ can dramatically increase with positive potentials, whilst the negative potentials increase the rate constants for switching the dye to the OFF state, *k*_*off*_. The values for *k*_*on*_ and *k*_*off*_ under electrochemical control is at least an order of magnitude greater than can be achieved optically. This is important as it means that electrochemical switching can give a much broader range of dye switching behavior depending on the imaging requirements of the experimental system being imaged which will endow EC-STORM with high optical resolution^44^. In conventional STORM, high emitter density could not be completely avoided when it comes to densely labelled biological samples. Supplementary Movie 6 applied cellular microtubule as a demonstration for this point. It showed that in conventional STORM, increase of 642 nm laser power did not decrease the emitter density but rather increased it, even though the *k*_*off*_ was expected to be increased. This is because the imaging laser has a dual effect that more Alexa 647 can be also activated from OFF state.^1, 45^ If we utilize EC-STORM in this case, a negative potential (−0.6 V) could reduce the emitter density to much lower level for accurate molecule localization. Being able to control the emitter density is important for image quality as will be further discussed below.

We next asked the question, what proportion of fluorophores could be recovered from the OFF state to the ON state using EC-STORM. A poly-L-lysine coated covered ITO surface with a very low density of Alexa 647 dyes adsorbed as single molecules (Supplementary Figure 9) was prepared. The number of molecules on the surface was quantified in oxygen scavenger buffer where there was no cysteamine so no switching of the Alexa 647 could occur. This allows us to determine that there was a coverage of 272 ± 28 Alexa 647 molecules in the imaging area of 25.6 × 25.6 μm^2^ (Fig. 2d). After adding cysteamine to the buffer, we did STORM imaging using both electrochemical or photochemical switching. With photochemical switching and using UV laser activation (as high as 200 W cm^-2^ of 405 nm laser intensity was applied) during the imaging period (2 min, consisting of 2000 frames of images) ∼65% were recovered (with a coverage of 178 ± 32 Alexa 647 molecules in the imaging area of ∼25.6 μm^2^, Fig. 2d-e). Importantly, with the electrochemical switching, 277 ± 39 Alexa 647 were counted per image area which represents 100% of Alexa 647 molecules in the OFF state being switched back to ON state. The results indicate EC-STORM shows higher recovery yield compared with conventional STORM, which might be owing to the minimization of photo-bleaching with the EC-STORM approach. This is because unlike conventional STORM, intense 642 nm laser illumination is not required for OFF-switching of the dyes for EC-STORM.

We next explored the utility of EC-STORM for cellular imaging by looking at microtubules in COS-7 cells (Fig. 3 a-c). The superior ability of EC-STORM, over conventional STORM, to precisely control the emitter density results in well-resolved tubulin structures under a negative potential (−0.3 V) without compromising the imaging speed. In comparison, under conventional STORM condition, part of microtubules in the highlighted region was not resolved as the overlapping emission prevents localization of the individual emitters (Fig. 3a, b, III vs. IV). We also assessed the STORM image resolution with Fourier ring correlation analysis^46^. A slightly improved resolution with EC-STORM over conventional STORM was observed, with estimated values from ∼53 nm to ∼41 nm (Fig. 3c, corresponding to STORM Fig. 3a I and EC-STORM Fig.3a II). The improvement in the image resolution with EC-STORM is more evident in the zoomed region where the tubulins are densely packed. Both the line profile (Fig. 3b) and the Fourier ring correlation curves (Fig. 3c, with measured resolution of ∼58 nm corresponding to Fig. 3a III *vs*. ∼45 nm corresponding to Fig. 3a IV) show the improved resolution of EC-STORM imaging.

**Fig. 3.**
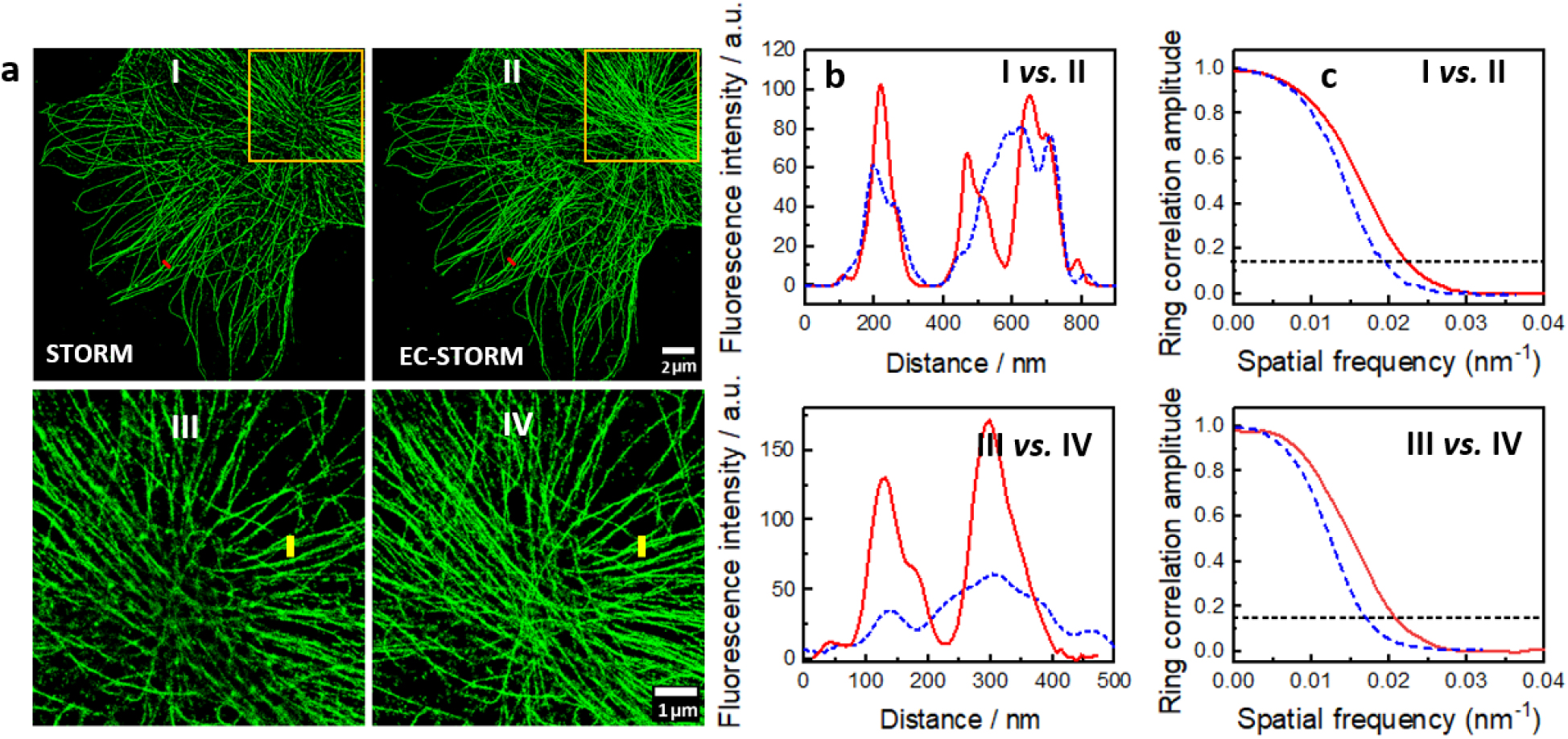
2D EC-STORM imaging. **a**, a comparison of 2D STORM images using conventional STORM (I and III) and EC-STORM (II and IV) methods at the same region of a cell. ‘STORM’ condition refers to 4 kW cm^-2^ 642 nm laser and no UV laser was applied, ‘EC-STORM’ condition refers to 4 kW cm^-2^ 642 nm laser and applying a potential at -0.3 V. The imaging COS-7 cells were labeled for microtubules with Alexa 647. Both the 2D STORM images were reconstructed from 15,000 frames of TIRF images, with camera exposure time of 33 ms. The zoomed region shows that the electrochemical potential can improve image resolution by decreasing the emitter density. **b**, Line profiles (blue dash for ‘STORM’, red line for ‘EC-STORM’) corresponding to the green and yellow marks in the left panel **a. c**, Fourier ring correlation curves corresponding to the left STORM image and zoomed region in panel **a**, blue line for ‘STORM’, red line for ‘EC-STORM’.

We also tested the EC-STORM for imaging more challenging specimens such as nuclear pore complexes and clathrin coated pits in COS-7 cells (Supplementary Figure 11 a, d), where the diameters of the structures are close to 100 nm. The octagonal distribution of Nup98 within nuclear pore complex and the spherical nature of the mature clathrin coated pit were both clearly revealed in EC-STORM. Our results demonstrated that EC-STORM showed comparable image quality with conventional STORM to resolve those cellular nanoscopic structures (Supplementary Figure 11 b-c, e-f). Nevertheless, during image acquisition EC-STORM offers more straightforward control over emitter density by directly changing the applied potential rather than balancing the thiol concentrations and intensities of UV laser in conventional STORM approach.

We further asked whether the EC-STORM could be used for 3D STORM imaging. To test the working depth from the surface using electrochemical activation, we imaged Alexa 647 labelled polystyrene beads with diameter of 100 μm, which confirms the electrochemical switching is applicable for at least 2 μm along the z-direction (Supplementary Movie 7 and Supplementary Figure 12). Then a 3D EC-STORM image (Supplementary Figure 13) of microtubules in COS-7 cells labelled with Alexa 647 revealing the z-dimension information (color-coded) was acquired. This experiment indicates electrochemical activation is feasible for 3D STORM imaging.

The key advantage of EC-STORM however goes beyond imaging towards molecular counting. The ability of STORM to image individual single fluorophores invokes the proposition of quantitative single molecule counting using localization microscopy^47-53^. However, molecular counting in conventional STORM can be quite difficult in practice. The uncertainty of labelling degree in immunostaining,^53^ inactivation or photobleaching of fluorophores resulting under-counting^53^, and random re-blinking induced over-counting^18, 47^ can all complicate the molecule quantification. As compared to the conventional STORM, EC-STORM shows promising features: it has 100% recovery yield to resolve all the fluorophores and it allows for reversible and fast fluorophore switching where the random fluorophore blinking could be possibly suppressed. Owing to these features, we designed a single molecule counting strategy using EC-STORM as illustrated in Fig. 4a. The electrochemical potential was pulsed between negative (−0.6 V) and positive (0.1 V) potentials for many cycles. With longer durations at the negative potential, the Alexa 647 is held in OFF state and will only get the change to be turned ON during the short positive potential pulse (Supplementary Movie 8). Results in Fig. 4b-c support our hypothesis that the random blinking of Alexa 647 was suppressed efficiently under negative potential, and over 95% of the ON events occur at positive potential pulse. To statistically study the blinking dynamics of Alexa 647 while pulsing the potential, we further performed an autocorrelation analysis^54^ of single Alexa 647 molecules in conventional STORM condition with UV laser activation and EC-STORM condition with pulsed potential activation. The autocorrelation time determined from the autocorrelation decay reveals the average ON time of Alexa 647,^55^ was significantly shorter for EC-STORM condition with pulsed potentials settings relative to conventional STORM. The decreased average ON time indicates the ON state Alexa 647, mostly appear under the positive potential pulse, and are turned to the OFF state by the followed negative potential pulse.

**Fig. 4.**
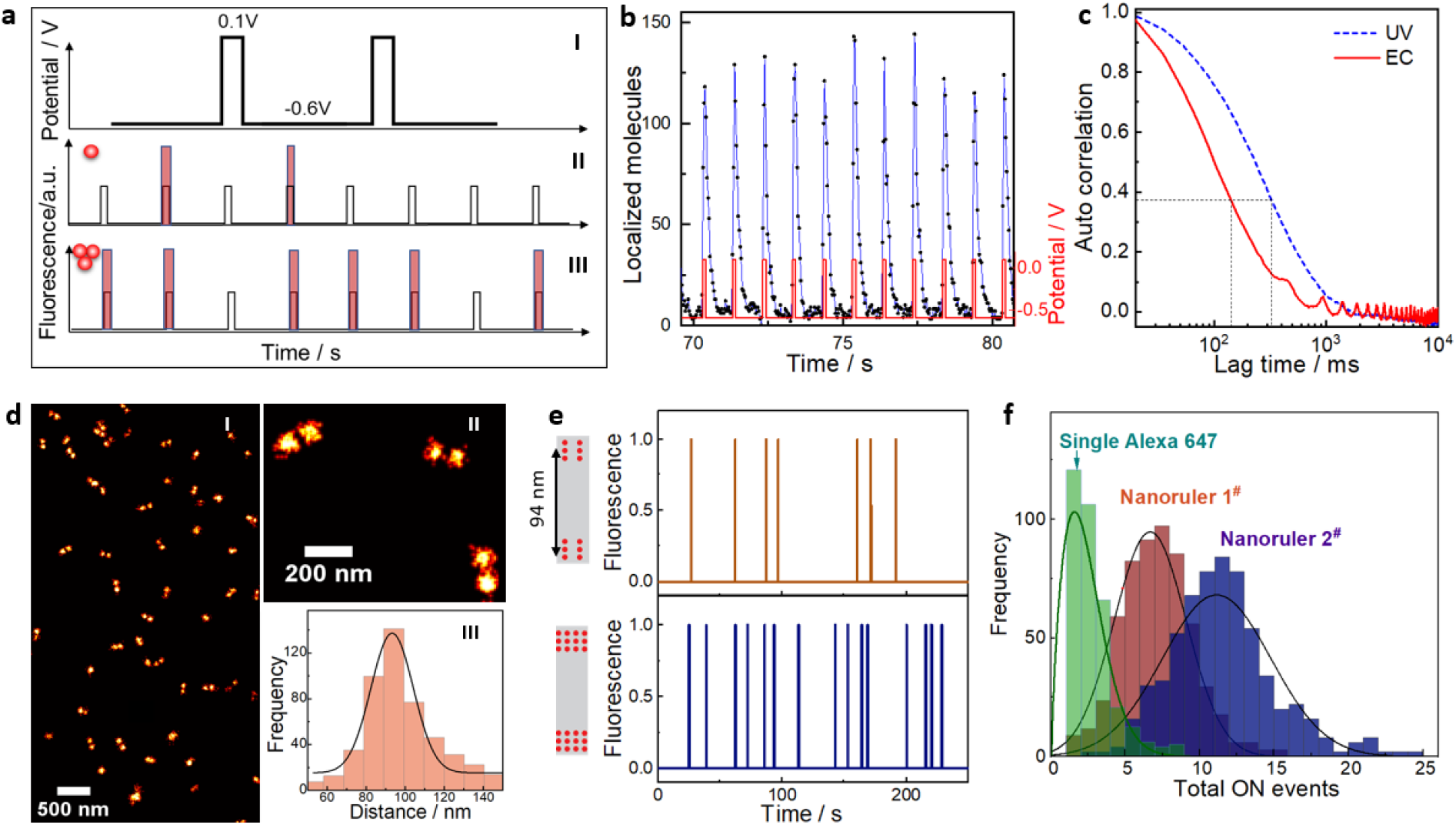
Single molecule counting exploration by EC-STORM. **a**, (I) Schematic of the pulsed electrochemical potential (−0.6 V for 900 ms and 0.1 V for 100 ms for multiple cycles) for electrochemical switching of the fluorophores. All the fluorophores could be switched off at -0.6 V, and only a small subset of the fluorophores would be switched on under a short pulse of 0.1 V potential, and then switched off again immediately after re-applying the negative potential. As illustrated in II-III, assuming the probability of each fluorophore being switched ON is identical, the sample with three fluorophores would exhibit three times more ON events than a single fluorophore. The red bar represents fluorescent ON events and the black line represents the underlying applied electrochemical potentials. **b**, The number of ON events (blue line) for Alexa 647 labeled DNA origamis in the field of view (25 × 25μm^2^) as a function of the pulsed electrochemical potential (red line). The fluorescence was acquired with 1 kW cm^-2^ of 642 nm laser excitation. **c**, Autocorrelation analysis of the fluorescent trajectory for Alexa 647 labeled DNA origami under conventional STORM conditions (blue dashed line, 1 kW cm^-2^ of 642 nm laser and 10 W cm^-2^ of UV laser were on) and EC-STORM where the potential was pulsed (red line, 1 kW cm^-2^ of 642 nm laser was on). The average correlation time at 1/e of the maximal autocorrelation amplitude were highlighted in dotted line, which was ∼323 ms and ∼138 ms for conventional STORM and EC-STORM (pulsed potential) respectively. **d**, EC-STORM image of DNA nanoruler acquired by reconstructing 5000 frames of images while pulsing the electrochemical potentials for 250 cycles. Two ends of the DNA nanoruler were labelled with Alexa 647 and were separated by 94 nm. **e**, Example for the fluorescent trajectories of two types of DNA nanorulers while pulsing the electrochemical potentials. Representative schematic of the two types of nanoruler is shown on the left, where the two ends of the DNA nanoruler had either 6 or 12 binding sites for Alexa 647 and that were separated by 94 nm. **f**, By analyzing thousands of fluorescent trajectories under pulsing electrochemical potentials, histograms were obtained for the total ON events counted for various DNA origami samples. The single Alexa 647 refers to the origami with single Alexa 647, nanoruler 1^#^ and 2^#^ refers to the DNA nanorulers illustrated in panel e with 6 or12 binding sites at each end.

If we assume the probability of switching ON a fluorophore is identical for each Alexa 647 at the positive potential pulse, the probability of switching ON events in each location is therefore determined by the total underlying number of Alexa 647 molecules in that location. With calibration from a known site, the number of Alexa 647 in unknown regions can then be calculated based on the ON events frequency in a given time interval. As proof of principle, we applied EC-STORM to count a DNA nanoruler sample designed with Alexa 647 labels at two ends which were separated by 94 nm. The STORM images (Fig. 4d) showed consistent observation with the expected fluorophore separation. Fig. 4e shows an example of the fluorescence trajectories of two types of DNA origami samples with 6 × 2 and 12 × 2 binding sites for Alexa 647, respectively. Much more ON events were observed for the nanoruler with 12× 2 binding sites. To exclude the random re-blinking events, only the ON events occurred at positive potential were analyzed. Histograms in Fig. 4f reflect the differences for the two groups of DNA nanorulers with 6 × 2 and 12 × 2 binding sites for Alexa 647, respectively. To quantify the absolute number of Alexa 647 on nanoruler, we used a DNA origami sample with only one Alexa 647 label for calibration. A histogram of total ON events for a single Alexa 647 under pulsed electrochemical potential shows the peak centered at 1.68 ± 0.10. From this calibration number of 1.68 ± 0.10 ON events per fluorophore under these conditions the number of Alexa 647 for the two types of DNA origami nanorulers were 4.10 ± 0.18 and 7.07 ± 0.29, respectively (Table 1). This quantification of Alexa 647 molecules indicated the labelling of the DNA origami was lower than the number of available binding sites due to incomplete labelling efficiency. When the ON events of the same nanorulers were counted under conventional STORM conditions (Supplementary Figure 14), there is no clear separation for the two group of nanorulers due to fluorophore re-blinking, and hence such quantification cannot be performed.

**Table 1.** Number of Alexa 647 counted by EC-STORM and electrochemical absolute counting.

| Sample | Number of Alexa 647 obtained from EC-STORM | Number of Alexa 647 obtained from maximum brightness |
| --- | --- | --- |
| Single Alexa 647 | 1 | 1 |
| Nanoruler 1<sup>#</sup> | 4.10 ± 0.18 | 4.04 ± 0.10 |
| Nanoruler 2<sup>#</sup> | 7.07 ± 0.29 | 7.24 ± 0.21 |

To evaluate the accuracy of the molecular counting with EC-STORM, we also estimate the number of Alexa 647 on DNA nanoruler by measure the maximum brightness of the nanorulers. In detail, the electrochemical potential was ramped from -0.8 V to 0.6 V that all Alexa 647 was switched to ON state at the high potential (Supplementary Figure 15). When the electrochemical potential was ramped back to -0.8 V, the ON state molecules were all turned OFF. This was repeated for three cycles. Subsequently, an EC-STORM image was obtained in the same field of view to locate each DNA origami. The spatial coordinates were used to extract the fluorescence trajectory during electrochemical potential ramping experiment and estimate the total Alexa 647 number at each origami from the maximum brightness. Our results showed that the average numbers of Alexa 647 of the complete resolved nanorulers with 6 × 2 and 12 × 2 binding sites were consistent with the results obtained from EC-qSTORM (Table 1).

## Conclusions

Single molecule localization based STORM technique demands well controlled ON and OFF switching of the organic dyes. Here we presented an electrochemical switching method referred to as EC-STORM that affords robust control of the switching using Alexa 647 dye as example. We showed the amount of Alexa 647 in its ON state could be controlled by the applied potential and suggested the switching is based on the electrochemical reversible thiol-ene reaction. With C-S covalent bond formation and dissociation at the polymethine bridge, the conjugated π-electron system of the Alexa 647 will be disrupted or restored which represent the OFF and ON state of the fluorophore respectively.

Our experiments show the EC-STORM measurements have several promising benefits over conventional STORM. Firstly, substituting electrochemical for photochemical switching eliminates the needs for initial high powered laser illumination to switch molecules to OFF state and ultraviolet laser to reactivate the molecules during data acquisition. As such, unwanted photodamage and photobleaching is dramatically mitigated. A second benefit is the fine control over the fluorophore state provided using electrochemical switching mean the number of fluorophores in the ON state can be precisely controlled such that regions with high densities of labels, which could not be feasibly imaged with conventional STORM can be imaged with EC-STORM. Similarly, if there are very few ON state Alexa 647 on the surface, applying a more positive electrochemical potential turns more fluorophores into the ON state such that 100% of the fluorophores on the surface can be imaged. A third advantage, the reversible and fast ON and OFF switching of Alexa 647 using electrochemical potential allows molecular counting. With all the fluorophores switching to OFF state by a negative potential, the probability of fluorescent ON events under a positive potential is determined by the underlying number of fluorophores. The working principle for molecular counting using EC-STORM is similar to the quantitative PAINT^56, 57^, while EC-STORM only requires standard dye labelled antibodies, reduces the high fluorescence background from floating imager strands, and shows faster imaging and counting process. We demonstrated the strategy using programmable pulsed electrochemical potential for molecule counting of Alexa647 on custom designed nanorulers. It is anticipated that EC-STORM can be used to perform molecular counting in biological samples as well.

Taken together, electrochemical switching of fluorophores to control their blinking behavior, has the potential to transform single molecule localization microscopy from a powerful imaging tool to a single molecule imaging tools with a broader range of imaging capabilities that allows quantitative molecular counting inside cells. Accompanying this dramatic increase in capabilities is the potential simplification of the microscopes to only requiring an imaging laser and an off-the-shelf electrochemical potentiostat.

## Methods

### Chemicals and materials

Alexa 647 labelled Goat anti-Rabbit IgG secondary antibody and anti-alpha tubulin primary antibody was purchased from Thermo Fisher Scientific (Australia). All solvents used were analytical grade unless further indicated. All chemicals, unless noted otherwise, were of analytical grade and used as received from Sigma-Aldrich (Australia). Aqueous solutions were prepared with Milli-Q water of 18.2□MΩ□cm resistivity.

Oxygen scavenging tris buffer was prepared in the following way: (1) stock buffer A containing 50 mM Tris, 10 mM NaCl (adjust to pH 8), (2) 10% glucose was added to buffer A, followed by adding 0.5 mg/mL glucose oxidase and 40 μg/mL catalase.

STORM buffer through the work was prepared by adding 50 mM cysteamine to the oxygen scavenging tris buffer. In Supplementary Figure 1a-b, the buffer solution was prepared by adding 2 mM Trolox to the oxygen scavenging tris buffer. In Supplementary Figure 1c-d, the buffer solution was prepared by adding 1 mM potassium ferricyanide into the oxygen scavenging tris buffer.

#### Single Alexa 647 fluorophores covered ITO sample

was prepared by immersing poly-L-lysine coated ITO into 100 pM Alexa 647 in PBS buffer for 10 mins. Single molecule photobleaching curves in PBS buffer were performed to analyse the step-wise intensity drop and the statistical analysis confirms over 99% of the Alexa 647 are single fluorophores absorbs on ITO (Supplementary Figure 10).

#### BSA-Alexa 647 covered ITO sample preparation

Alexa Fluor 647 Protein Labelling Kit (Thermo Fisher Scientific, Australia) were used to label BSA (Sigma-Aldrich, United States), with a determined degree of labelling 2.46 (measured by Thermo Scientific™ NanoDrop™ One Microvolume UV-Vis Spectrophotometer, see the corresponding UV-Vis absorbance spectrum in Figure 11a). BSA-Alexa 647 solution (2 mg/mL) was diluted in blocking buffer containing with unlabelled BSA in PBS (5 mg/mL) with different dilution ratios (1:100, 1:200, 1:400, and 1:2000), and placed onto clean ITO surface, after physical absorption for 30 min and washing with PBS for three times, the surfaces were imaged in STORM buffer.

#### DNA origami covered ITO sample preparation

Immobilization of DNA origami (GATTAquant, Germany) was achieved by modifying a clean 22×22 cm ITO following the below protocols: Wash ITO three times with PBS and incubate the ITO with 200 μL of BSA-biotin solution (1 mg/mL in PBS) for 30 min. Remove the BSA-biotin solution and wash it three times with PBS. Make sure not to scratch with the pipette tip on the surface. Incubate the ITO with 200 μL of neutravidin solution (1 mg/mL in PBS) for 30 min. Remove the neutravidin solution and wash the chamber three times with 1x PBS supplemented with 10 mM magnesium chloride (so-called immobilization buffer). Dilute 2 μL of the DNA origami solution with 200 μL immobilization buffer. Incubate the ITO with this solution for 30 min. Wash the ITO three times with immobilization buffer and stored in the immobilization buffer for imaging.

#### Immunofluorescence staining

Immunostaining was performed using COS-7 cells (ATCC CRL-1651) cultured with Dulbecco′s Modified Eagle′s Medium fortified with 10% FBS, penicillin and streptomycin, and incubated at 37°C with 5% CO_2_. COS-7 cells were plated in 6-well plates with ITO slides in at ∼10,000-20,000 cells per well on the day before fixation. 10 ul of gold nanoroad (A12-25-650-PAA-DUG-25 from NanoParTz) were added to the ITO coverslip for fiducial drift correction.

The immunostaining procedure for microtubules consisted of: fixation for 7 min at 37°C with 4% paraformaldehyde (Sigma) in PBS, washing with PBS, permeabilization for 5 min with 0.2% Triton X-100 and 3% BSA in PBS, incubation for 1.5 h with rabbit anti α-tubulin monoclonal antibody (ab216650, Abcam) diluted to 2 μg mL^−1^ in blocking buffer (0.2% Triton X-100 and 3% BSA in PBS), washing with PBS, incubation for 30 min with secondary anti-Rabbit IgG antibodies labelled with Alexa 647 at a concentration of ∼2.5 μg mL^−1^ in PBS; washing with PBS, fixation for 5 min with 4% paraformaldehyde in PBS and finally washing with PBS.

The immunostaining procedure for nuclear pore complex and clathrin-coated pits consisted of: fixation for 20 min at -20°C with 100% methanol, rinsing with PBS, permeabilization for 5 min in 0.1% w/v saponin in PBS, incubation for 6 h with primary antibody of anti Nup98 mAb 2H10 (ab50610, Abcam) or anti clathrin Heavy Chain (MA1-065, Thermo Fisher) at 1 μg mL^−1^ in PBS blocking buffer with 5% BSA and 0.01 % w/v saponin, washing with PBS containing 0.5 % Tween-20 and 0.01% saponin, incubation for 1h with Alexa 647 conjugated secondary antibodies of goat against rat IgG (ab 150159, Abcam) or goat anti mouse IgG (A-21235, Thermos Fisher) at a concentration of 2 μg mL^−1^ in PBS with 2.5% BSA and 0.01% saponin, fixation for 10 min with 4% paraformaldehyde in PBS and finally washing with PBS.

#### Electrochemistry

Cyclic voltammetry, chronoamperometry, and different pulsed voltammetry were performed by SP-200 Potentiostat (Bio-Logic, France). All the electrochemistry was carried out in a custom chamber (Chamlide EC 22, Live Cell Instrument Co., Ltd., Republic of Korea) containing an Ag|AgCl|3M KCl reference electrode and a Pt-wire counter-electrode, where the working electrode was the indium tin oxide (ITO) coated coverslips (8-12 Ω, 22×22 cm, SPI Supplies, USA).

#### STORM imaging and analysis

The TIRF images were collected on Zeiss Elyra SP2 Super-resolution PALM microscope. The collimated and linearly p-polarized 642 nm laser was reflected from the 642 nm long pass dichroic mirror and focused at the back focal plane of the 100 X 1.46 NA Oil objective. The focus is laterally shifted alone the back focal plane to provide either EPI (0°) or TIRF (66.7°) illumination. The TIRF angle was identical between the glass and ITO surface. The fluorescence was collected by the same objective and guided to a cooled electron multiplying charge-coupled Device EMCCD camera (iXon DU-897). For STORM imaging, 2-4 kW cm^-2^ 642 nm laser was used for excitation with less than 50 W cm^-2^ 405 nm laser for photoactivation. For electrochemistry-based STORM, only 1-4 kW cm^-2^ 642 nm laser is used. For 2D STORM, 2,000-20,000 images were acquired on a single EMCCD camera with an exposure time of 33-100 ms and camera gain of 80-120. Either Cross-correlation based drift correction that provided in Zeiss Zen software or the gold nanorods contained in the image were used for fiducial based drift correction. When reconstructing 2D STORM, localizations were filtered by the localization precision of less than 35□nm.

For 3D STORM, 30,000 images were acquired simultaneously on two separate EMCCD cameras with exposure time of 33 ms. The exposure time of the two cameras was synchronized by the internal trigger provided in the Zeiss Elyra. The Alexa 647 fluorescence was split evenly onto two cameras by a long pass dichroic beam splitter (690 nm, AHF Analysetechnik). The fluorescence was focused onto individual cameras by two independent motorized tube lenses. The tube lens in second light path was moved closer to the camera chip so that molecules located at 900 nm higher position in the sample space become focused on this camera. During 3D acquisition, the objective was parked at the middle of the two focal planes to produce maximal contrast in the shape variation of the PSF across the two cameras. Prior to 3D image acquisition, the 100 nm microspheres (Orange fluorescent (540/560 nm), Thermo Fisher Scientific) sample was imaged in TIRF mode for two camera alignments. The mirror prior to the tube lens in the second light path were adjusted so that the bead images in two cameras were maximally overlayed in xy direction. For 3D calibration the piezo stage was moved up at 50 nm step size from -1350 nm to +1350 nm relative to the middle point of two focal plane. A molecular density filter of > 5 molecules were applied in the reconstructed STORM image to remove sparse background due to nonspecific binding of antibodies.

#### Rate constant for Alexa 647 switching

To measure the rate constant *k*_*on*_ and *k*_*off*_ using electrochemical switching method, the Alexa 647 labelled tubulin samples were imaged under 642 nm reading laser (3 kW cm^-2^). By applying a negative potential to the ITO surface, Alexa 647 will be switched into the OFF state, followed a positive potential will reversibly switch the Alexa 647 back to ON state. The rate constant was determined by counting the number of molecules in the field of view as function of time and then fitting the distribution to a single exponential function (Supplementary Figure 9). For comparison, the rate constant for *k*_*on*_ and *k*_*off*_ using photochemical switching was measured by irradiated the sample with strong 642 nm laser to switch Alexa 647 off (*k*_*off*_) first, and then reduce the 642 nm laser to 3 kW cm^-2^ on and turn UV light on to switch the OFF state fluorophores back to ON state (*k*_*on*_).

#### Recovery yield comparison

The STORM images were obtained under different conditions at the same single Alexa-647 molecule covered ITO sample but at different regions. 2,000 frames images were collected to reconstruct the final STORM image. ‘No switching’ represents the STORM image in oxygen scavenger buffer without cysteamine that all the Alexa 647 molecules should be in the ON state. Under ‘EC’ condition, a -0.6 V potential was applied for 30 s for off switching the dyes and then a positive potential was increased from -0.1 to 0.35 V during data collection to ensure as more as Alexa 647 could be activated to ON state. Under ‘UV’ condition, a 10 kW cm^-2^ of 642 nm laser was applied for 30 s for initial off-switching the dyes, and the UV laser intensity was increased from 10 to 200 W cm^-^ during data collection. A 2 kW cm^-2^ of 642 nm laser was on during data collection for all the conditions. Under ‘EC’ and ‘UV’ conditions, after 500 frames, there are very few molecules to make sure we have imaged nearly all the molecules that have been turned on.

#### DNA origami imaging and single molecule counting

For STORM imaging of DNA origami, 6,000 images were acquired with an exposure time of 50 ms and camera gain of 100. To filter out the complete nanorulers with two labelling sites, we first examined the pairwise Euclidean distance between all possible pairs of localization events in the STORM table. Localization events that belong to the same nanoruler tends to be more closely packed in space, therefore, the pairwise distance between the localization events of the same nanoruler was much smaller than distance between different nanorulers. Once the densely packed, nanoruler like regions were isolated. We analysed the distance between localization events inside the region of interest to its centre of the mass. A single spotted nanoruler due to misfolding related damage would produce a much smaller value compared to a dual spotted nanoruler, and hence will be excluded from the STORM image. Once the dual spotted nanoruler were identified (as shown in Fig. 4a), the localization events within the nanoruler were segregated into two groups based on the pairwise molecule distance. The distance between these two groups (from centre to another centre of the group) will be analysed (Fig.4b).

Once the localization events of fluorophores within each spot of the nanoruler were identified. The temporal information of the localization events was examined to regroup the self-blinking induced repeating localizations. Here, we apply a hash threshold of 6 seconds based on the previous knowledge that for 99.9% of Alexa 647, the blinking of the same molecule occurred with short intervals of less than 6 seconds. The assumption is that if the temporal difference between the localization evens within the same spot is shorter than 6 seconds, it is counted as one molecule. If the localization events were more than 6 second apart, they will be counted as two molecules. Using this method, we counted at each end of the nanoruler there are around 4 Alexa 647 molecule under UV activation mode, and around 3 Alexa 647 under electrochemical activation.

#### Molecular counting using pulsed electrochemical potential (EC-qSTORM)

The electrochemical potential was pulsed between -0.6 V for 900 ms and 0.1 V for 100 ms for 250 cycles. A STORM image was generated while the potential was pulsing. From the constructed STORM image we can located each DNA origami, we then analyze the fluorescence trajectories of these well-resolved DNA nanoruler and count for the total ON events.

#### Absolute molecule counting at DNA origami using electrochemical switching method

We ramped the electrochemical potential between -0.8 V and 0.6 V to sequentially turn on and off all the Alexa 647 at each DNA origami. A stepwise increase or decrease of the fluorescence intensity indicates the molecule transfer between ON and OFF states were guided by the potential. To increase the statistics of the molecule counting, we repeated the potential ramping for three cycles. Followed, a STORM imaging was performed at the same field of view. 5,000 images were acquired with an exposure time of 50 ms. The electrochemical potential was slowly increased from 0-0.4 V during image acquisition. From the reconstructed STORM image, we can locate each DNA origami. We then analyzed the fluorescence trajectory in electrochemical potential ramping experiment and counted the total Alexa 647 number at each origami from the maximum brightness.

## Data availability

Source data are provided with this paper at:

https://doi.org/10.5061/dryad.7pvmcvdx9

Part of the raw data for EC-STORM or conventional STORM are not suitable for distribution through public repositories due to the large file size and are available from the corresponding author upon request.

## Code availability

Custom MATLAB (MathWorks) codes used for data processing are available at: https://github.com/mayuanqing8/EC-STORM

## Acknowledgments

We thank S. Ciampi, T.□ Boecking for comments and discussions, and G. Ball for sharing insights with the experimental system. We acknowledge technical assistant by the Katharina Gaus Light Microscopy Facility, University of New South Wales.

## Author contributions

J.J.G. and K.G. designed the project. Y.Y. and Y.M. conceived the experiments, analyzed the data, and wrote the manuscript. Y.M. wrote the custom code in MATLAB for analyzing the data. R.T. helped revise the manuscript. J.J.G supervised all the work and wrote the manuscript.

## Supplementary information

Supplementary Figures 1 to 15 and supplementary movies 1-8 are available in the supplementary information.

## References

1. Huang, B., Bates, M. & Zhuang, X. Super-resolution fluorescence microscopy. Annu. Rev. Biochem. 78, 993–1016 (2009).

2. Hell, S.W. & Wichmann, J. Breaking the diffraction resolution limit by stimulated emission: stimulated-emission-depletion fluorescence microscopy. Opt. Lett. 19, 780–782 (1994).

3. Klar, T.A., Jakobs, S., Dyba, M., Egner, A. & Hell, S.W. Fluorescence microscopy with diffraction resolution barrier broken by stimulated emission. Proc. Natl. Acad. Sci. 97, 8206–8210 (2000).

4. Dertinger, T., Colyer, R., Iyer, G., Weiss, S. & Enderlein, J. Fast, background-free, 3D super-resolution optical fluctuation imaging (SOFI). Proc. Natl. Acad. Sci. 106, 22287–22292 (2009).

5. Fernández-Suárez, M. & Ting, A.Y. Fluorescent probes for super-resolution imaging in living cells. Nat. Rev. Mol. Cell Biol. 9, 929–943 (2008).

6. Shroff, H., Galbraith, C.G., Galbraith, J.A. & Betzig, E. Live-cell photoactivated localization microscopy of nanoscale adhesion dynamics. Nat. Methods 5, 417–423 (2008).

7. Jones, S.A., Shim, S.-H., He, J. & Zhuang, X. Fast, three-dimensional super-resolution imaging of live cells. Nat. Methods 8, 499–505 (2011).

8. Sengupta, P., Jovanovic-Talisman, T., Skoko, D., Renz, M., Veatch, S.L., Lippincott-Schwartz, J. Probing protein heterogeneity in the plasma membrane using PALM and pair correlation analysis. Nat. Methods 8, 969–975 (2011).

9. Sengupta, P., Jovanovic-Talisman, T. & Lippincott-Schwartz, J. Quantifying spatial organization in point-localization superresolution images using pair correlation analysis. Nat. Protoc. 8, 345–354 (2013).

10. Rust, M.J., Bates, M. & Zhuang, X. Sub-diffraction-limit imaging by stochastic optical reconstruction microscopy (STORM). Nat. Methods 3, 793–796 (2006).

11. Betzig, E., Patterson, G.H., Sougrat, R., Lindwasser, O.W., Olenych, S., Bonifacino, J.S., Davidson, M.W., Lippincott-Schwartz, J., Hess, H.F. Imaging intracellular fluorescent proteins at nanometer resolution. Science 313, 1642–1645 (2006).

12. Hess, S.T., Girirajan, T.P. & Mason, M.D. Ultra-high resolution imaging by fluorescence photoactivation localization microscopy. Biophys. J. 91, 4258–4272 (2006).

13. Sharonov, A. & Hochstrasser, R.M. Wide-field subdiffraction imaging by accumulated binding of diffusing probes. Proc. Natl. Acad. Sci. 103, 18911–18916 (2006).

14. Betzig, E. Proposed method for molecular optical imaging. Opt. Lett. 20, 237–239 (1995).

15. Heilemann, M., Van De Linde, S., Schüttpelz, M., Kasper, R., Seefeldt, B., Mukherjee, A., Tinnefeld, P., Sauer, M. Subdiffraction-resolution fluorescence imaging with conventional fluorescent probes. Angew. Chem., Int. Ed. 47, 6172–6176 (2008).

16. Endesfelder, U. & Heilemann, M. Art and artifacts in single-molecule localization microscopy: beyond attractive images. Nat. Methods 11, 235–238 (2014).

17. Lelek, M., Gyparaki, M.T., Beliu, G., Schueder, F., Griffié, J., Manley, S., Jungmann, R., Sauer, M., Lakadamyali, M., Zimmer, C. Single-molecule localization microscopy. Nat. Rev. Methods Primers 1, 1–27 (2021).

18. Zhao, Z.W., Roy, R., Gebhardt, J.C.M., Suter, D.M., Chapman, A.R., Xie, X.S. Spatial organization of RNA polymerase II inside a mammalian cell nucleus revealed by reflected light-sheet superresolution microscopy. Proc. Natl. Acad. Sci. 111, 681–686 (2014).

19. Hilczer, M., Traytak, S., Tachiya, M. Electric field effects on fluorescence quenching due to electron transfer. J. Chem. Phys. 115, 11249–11253 (2001).

20. Guille-Collignon, M., Delacotte, J., Lemaître, F., Labbé, E. & Buriez, O. Electrochemical Fluorescence Switch of Organic Fluorescent or Fluorogenic Molecules. Chem. Rec. 21, 2193–2202 (2021).

21. Djoumer, R., Chovin, A., Demaille, C., Dejous, C. & Hallil, H. Real-time Conversion of Electrochemical Currents into Fluorescence Signals Using 8-Hydroxypyrene-1, 3, 6-trisulfonic Acid (HPTS) and Amplex Red as Fluorogenic Reporters. ChemElectroChem 8, 2298–2307 (2021).

22. Fan, S., Webb, J.E., Yang, Y., Nieves, D.J., Gonçales, V.R., Tran, J., Hilzenrat, G., Kahram, M., Tilley, R.D., Gaus, K., Gooding, J.J. Observing the Reversible Single Molecule Electrochemistry of Alexa Fluor 647 Dyes by Total Internal Reflection Fluorescence Microscopy. Angew. Chem., Int. Ed. 131, 14637–14640 (2019).

23. Li, H. & Vaughan, J.C. Switchable fluorophores for single-molecule localization microscopy. Chem. Rev. 118, 9412–9454 (2018).

24. Bates, M., Huang, B., Dempsey, G.T. & Zhuang, X. Multicolor super-resolution imaging with photo-switchable fluorescent probes. Science 317, 1749–1753 (2007).

25. Van de Linde, S. & Sauer, M. How to switch a fluorophore: from undesired blinking to controlled photoswitching. Chem. Soc. Rev. 43, 1076–1087 (2014).

26. Gidi, Y., Payne, L., Glembockyte, V., Michie, M.S., Schnermann, M.J., Cosa, G. Unifying Mechanism for Thiol-Induced Photoswitching and Photostability of Cyanine Dyes. J. Am. Chem. Soc. 142, 12681–12689 (2020).

27. Dempsey, G.T., Bates, M., Kowtoniuk, W.E., Liu, D.R., Tsien, R.Y. and Zhuang, X. Photoswitching mechanism of cyanine dyes. J. Am. Chem. Soc. 131, 18192–18193 (2009).

28. Vaughan, J.C., Dempsey, G.T., Sun, E. & Zhuang, X. Phosphine quenching of cyanine dyes as a versatile tool for fluorescence microscopy. J. Am. Chem. Soc. 135, 1197–1200 (2013).

29. Chibisov, A. Triplet states of cyanine dyes and reactions of electron transfer with their participation. J. Photochem. 6, 199–214 (1976).

30. van de Linde, S., Krstić, I., Prisner, T., Doose, S., Heilemann, M. and Sauer, M. Photoinduced formation of reversible dye radicals and their impact on super-resolution imaging. Photochem. Photobiol. Sci. 10, 499–506 (2011).

31. Baumann, R.P., Penketh, P.G., Seow, H.A., Shyam, K. & Sartorelli, A.C. Generation of oxygen deficiency in cell culture using a two-enzyme system to evaluate agents targeting hypoxic tumor cells. Radiat. Res. 170, 651–660 (2008).

32. Lu, X., Nicovich, P.R., Gaus, K. & Gooding, J.J. Towards single molecule biosensors using super-resolution fluorescence microscopy. Biosens. Bioelectron. 93, 1–8 (2017).

33. Lu, X., Nicovich, P.R., Zhao, M., Nieves, D.J., Mollazade, M., Vivekchand, S.R.C., Gaus, K., Gooding, J.J. Monolayer surface chemistry enables 2-colour single molecule localisation microscopy of adhesive ligands and adhesion proteins. Nat. Commun. 9, 1–10 (2018).

34. Parviz, M., Gaus, K. & Gooding, J.J. Simultaneous impedance spectroscopy and fluorescence microscopy for the real-time monitoring of the response of cells to drugs. Chem. Sci. 8, 1831–1840 (2017).

35. Koppenol, W.H., Stanbury, D.M. & Bounds, P.L. Electrode potentials of partially reduced oxygen species, from dioxygen to water. Free Radic. Biol. Med 49, 317–322 (2010).

36. Noël, J.-M., Latus, A., Lagrost, C., Volanschi, E. & Hapiot, P. Evidence for OH radical production during electrocatalysis of oxygen reduction on Pt surfaces: consequences and application. J. Am. Chem. Soc. 134, 2835–2841 (2012).

37. Anderson, L.C., Xu, M., Mooney, C.E., Rosynek, M.P. & Lunsford, J.H. Hydroxyl radical formation during the reaction of oxygen with methane or water over basic lanthanide oxide catalysts. J. Am. Chem. Soc. 115, 6322–6326 (1993).

38. Anbar, M., Meyerstein, D. & Neta, P. The reactivity of aromatic compounds toward hydroxyl radicals. J. Phys. Chem. 70, 2660–2662 (1966).

39. Neta, P. & Dorfman, L.M. in Pulse radiolysis studies. XIII. Rate constants for the reaction of hydroxyl radicals with aromatic compounds in aqueous solutions (ACS Publications, 1968).

40. Nordberg, J. & Arnér, E.S. Reactive oxygen species, antioxidants, and the mammalian thioredoxin system. Free Radic. Biol. Med 31, 1287–1312 (2001).

41. Fava, A., Reichenbach, G. & Peron, U. Kinetics of the thiol-disulfide exchange. II. Oxygen-promoted free-radical exchange between aromatic thiols and disulfides. J. Am. Chem. Soc. 89, 6696–6700 (1967).

42. Antonello, S., Benassi, R., Gavioli, G., Taddei, F. & Maran, F. Theoretical and electrochemical analysis of dissociative electron transfers proceeding through formation of loose radical anion species: reduction of symmetrical and unsymmetrical disulfides. J. Am. Chem. Soc. 124, 7529–7538 (2002).

43. Vaughan, J.C., Jia, S. & Zhuang, X. Ultrabright photoactivatable fluorophores created by reductive caging. Nat. Methods 9, 1181–1184 (2012).

44. Dempsey, G.T., Vaughan, J.C., Chen, K.H., Bates, M. & Zhuang, X. Evaluation of fluorophores for optimal performance in localization-based super-resolution imaging. Nat. Methods 8, 1027–1036 (2011).

45. Huang, B., Babcock, H. & Zhuang, X. Breaking the diffraction barrier: super-resolution imaging of cells. Cell 143, 1047–1058 (2010).

46. Banterle, N., Bui, K.H., Lemke, E.A. & Beck, M. Fourier ring correlation as a resolution criterion for super-resolution microscopy. J. Struct. Biol. 183, 363–367 (2013).

47. Annibale, P., Vanni, S., Scarselli, M., Rothlisberger, U. & Radenovic, A. Identification of clustering artifacts in photoactivated localization microscopy. Nat. Methods 8, 527–528 (2011).

48. Nicovich, P.R., Owen, D.M. & Gaus, K. Turning single-molecule localization microscopy into a quantitative bioanalytical tool. Nat. Protoc. 12, 453–460 (2017).

49. Durisic, N., Cuervo, L.L. & Lakadamyali, M. Quantitative super-resolution microscopy: pitfalls and strategies for image analysis. Curr. Opin. Chem. Biol. 20, 22–28 (2014).

50. Löschberger, A., Franke, C., Krohne, G., van de Linde, S. & Sauer, M. Correlative superresolution fluorescence and electron microscopy of the nuclear pore complex with molecular resolution. J. Cell Sci. 127, 4351–4355 (2014).

51. Wu, Y.-L., Tschanz, A., Krupnik, L. & Ries, J. Quantitative data analysis in singlemolecule localization microscopy. Trends Cell Biol. 30, 837–851 (2020).

52. Ehmann, N., Van De Linde, S., Alon, A., Ljaschenko, D., Keung, X. Z., Holm, T., Kittel, R. J. Quantitative super-resolution imaging of Bruchpilot distinguishes active zone states. Nat. Commun. 5, 4650 (2014).

53. Lee, S.-H., Shin, J.Y., Lee, A. & Bustamante, C. Counting single photoactivatable fluorescent molecules by photoactivated localization microscopy (PALM). Proc. Natl. Acad. Sci. 109, 17436–17441 (2012).

54. Basché, T., Moerner, W., Orrit, M. & Wild, U. Single-molecule optical detection, imaging and spectroscopy. (John Wiley & Sons, 2008).

55. Dickson, R.M., Cubitt, A.B., Tsien, R.Y. & Moerner, W.E. On/off blinking and switching behaviour of single molecules of green fluorescent protein. Nature 388, 355–358 (1997).

56. Jungmann, R.,Vendaño, M.S., Dai, M., Woehrstein, J.B., Agasti, S.S., Feiger, Z., Rodal, A., Yin, P. Nat. Methods 13, 439–442 (2016).

57. Strauss, S., Nickels, P.C., Strauss, M.T., Jimenez Sabinina, V., Ellenberg, J., Carter, J.D., Gupta, S., Janjic, N., Jungmann, R. Modified aptamers enable quantitative sub-10-nm cellular DNA-PAINT imaging. Nat. Methods 15, 685–688 (2018).

