## Supplementary Information for "Electrochemically controlled blinking of fluorophores to enable quantitative stochastic optical reconstruction microscopy (STORM) imaging"

**Supplementary Text**

**The fluorescence intensity of Alexa 647 can be tuned using the electrochemical potential.** Here we plated and fixed COS-7 cells stained with Alexa 647 labelled microtubules (α-tubulin) onto an ITO surface and imaged in an oxygen scavenger tris buffer (see Materials and Methods) with Trolox (Supplementary Figure 1a-b) or ferricyanide (Supplementary Figure 1c-d) as a redox mediator. TIRF images shows the microtubules are readily visible from the fluorescence of Alexa 647 but that the fluorescence intensity of Alexa 647 decreased at negative potentials (-0.7 V). Especially when ferricyanide is present, the fluorescence intensity of Alexa 647 dimmed to close to the background level at -0.7 V and could become bright again if the potential was returned to positive potentials (+0.7 V). The Supplementary Movie 1 shows the dynamic switching behaviour of Alexa 647 as the potential is cycled. Similarly, imaging in the oxygen scavenger buffer without ferricyanide or Trolox, the fluorescence intensity of Alexa 647 was almost unchanged as a function of applied electrochemical potential (see Supplementary Figure 2).

When Alexa 647 is in the ON state, a quaternary ammonium cation and a tertiary amine are linked by the conjugated polymethine bridge. The resonance between the two nitrogen atoms is essential for the fluorescence intensity of Alexa 647(*1*). Previous work has shown using Trolox as a redox mediator, that redox reactions can take place at the quaternary ammonium cation and alter the fluorescence behaviour of Alexa 647, which suggests the fluorescence intensity of Alexa 647 can be tuned using electrochemical potential(*2*). In this work we further show that using ferricyanide as a redox mediator, the fluorescence intensity of Alexa 647 can be more efficiently altered using the electrochemical potential (Fig. 1c and Supplementary Figure 2). All of the Alexa 647 dyes proceed to the same oxidation or reduction direction at applied positive or negative potential. Therefore, from the Supplementary Movie 1 we can see that by switching the potential between a negative and a positive potential, the bulk fluorescence intensity is becoming very dim and bright, respectively. This property might be applicable in other imaging methods such as structured illumination microscopy(*3, 4*).

**EC-STORM can better resolve underlying structure than conventional STORM.** After learning that the recovery yield for single Alexa 647 molecules using EC-STORM is higher than using the conventional UV activation pathway (Fig. 2d-e). We next asked the question as to whether the ability to completely recover all the OFF state Alexa 647 fluorophores with EC-STORM will give better resolve power of the underlying labelled structure than that which can be achieved using the conventional STORM approach. To answer this question we designed a surface covered with Alexa 647 labelled BSA with very low coverage (<1 BSA per um^2^) but where each BSA molecules was labelled with ~2-3 fluorophores. During data acquisition, we found that the electrochemical method could activate more Alexa 647 molecules than using UV laser (Supplementary Figure 10b). By analyzing the STORM images, we show that about twice as many BSA molecules were imaged using the positive potentials than when using laser switching in a given time. This finding indicates that even though there are more than one fluorophore labelled on one BSA molecule, photochemical activation could still not fully recover all the BSA. Hence part of BSA-Alexa 647 on ITO were still missed in the final reconstructed STORM images (Supplementary Figure 11c). The electrochemical activation has ~100% recovery capability for Alexa 647, and hence can better resolve the underlying BSA molecules.

**Supplementary Figures**

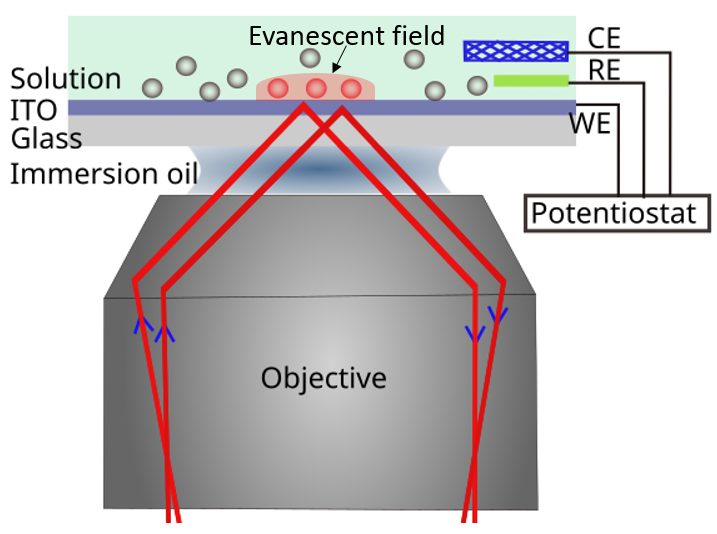

**Supplementary Figure 1**. Schematic illustration of the total internal reflectance (TIRF) microscope integrated with the electrochemistry set-up. A typical three-electrode system is utilized, which includes a transparent ITO covered glass slide as the working electrode (WE), a silver/silver chloride wire in 3 M KCl as the reference electrode (RE), and a platinum mesh counter electrode. Each electrode is connected to a potentiostat, which applies a defined potential difference between the WE and RE and monitor the current flow between the WE and CE. In TIRF microscopy the laser is totally internally reflected at the ITO/ water interface, and an evanescent field is created such that only fluorophores in the thin layer close to ITO surface will be excited. The red colored spheres indicate excited fluorophores, and the grey spheres imply unexcited fluorophores.

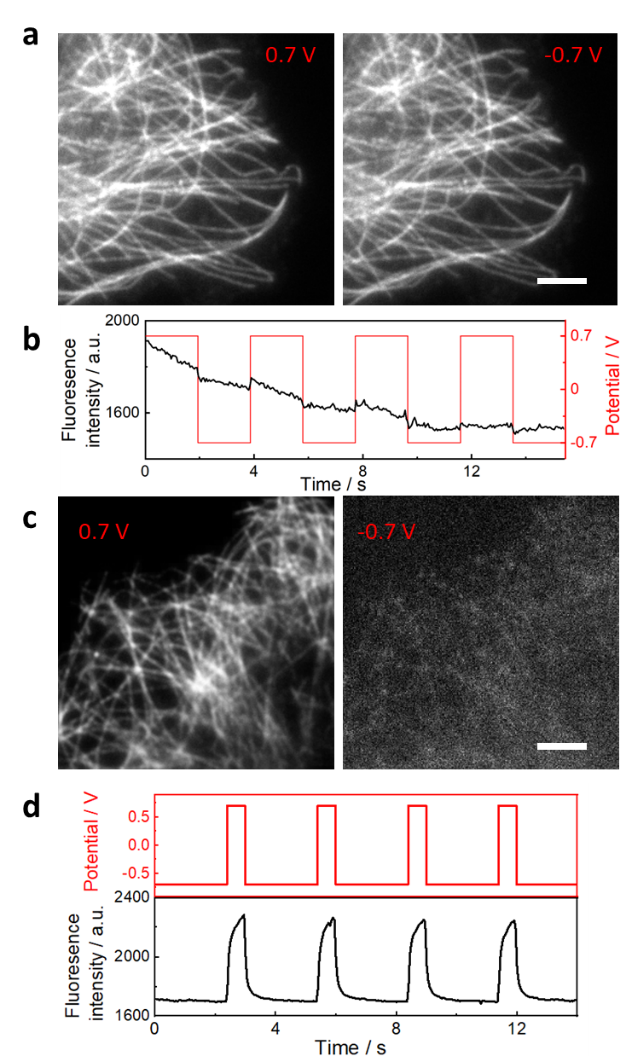

**Supplementary Figure 2. a,** TIRF images of Alexa 647 labelled microtubules (α-tubulin) of COS-7 cell in oxygen scavenger tris buffer with 2 mM Trolox at +0.7 V (left) and -0.7 V (right). All the Alexa 647 stays in the bright state while the overall fluorescence intensity is slightly higher at 0.7 V than the values at -0.7 V. The decay in fluorescence intensity was attributed to photobleaching during the images acquisition, **b**, the corresponding electrochemical potential modulated fluorescence intensity change curves for a. **c**, TIRF images of Alexa 647 labelled microtubules (α-tubulin) of COS-7 cell in oxygen scavenger tris buffer with 1 mM potassium ferricyanide at +0.7 V (left) and -0.7 V (right). All of the Alexa 647 are in bright state at positive potential (0.7 V) and are switched to a dim state at negative potential (-0.7 V), **d**, the corresponding electrochemical potential modulated fluorescence intensity change curves for c. A 642 nm laser with intensity of 0.1 kW cm^-2^ was used as the excitation laser for the imaging. The scale bar is 5 μm.

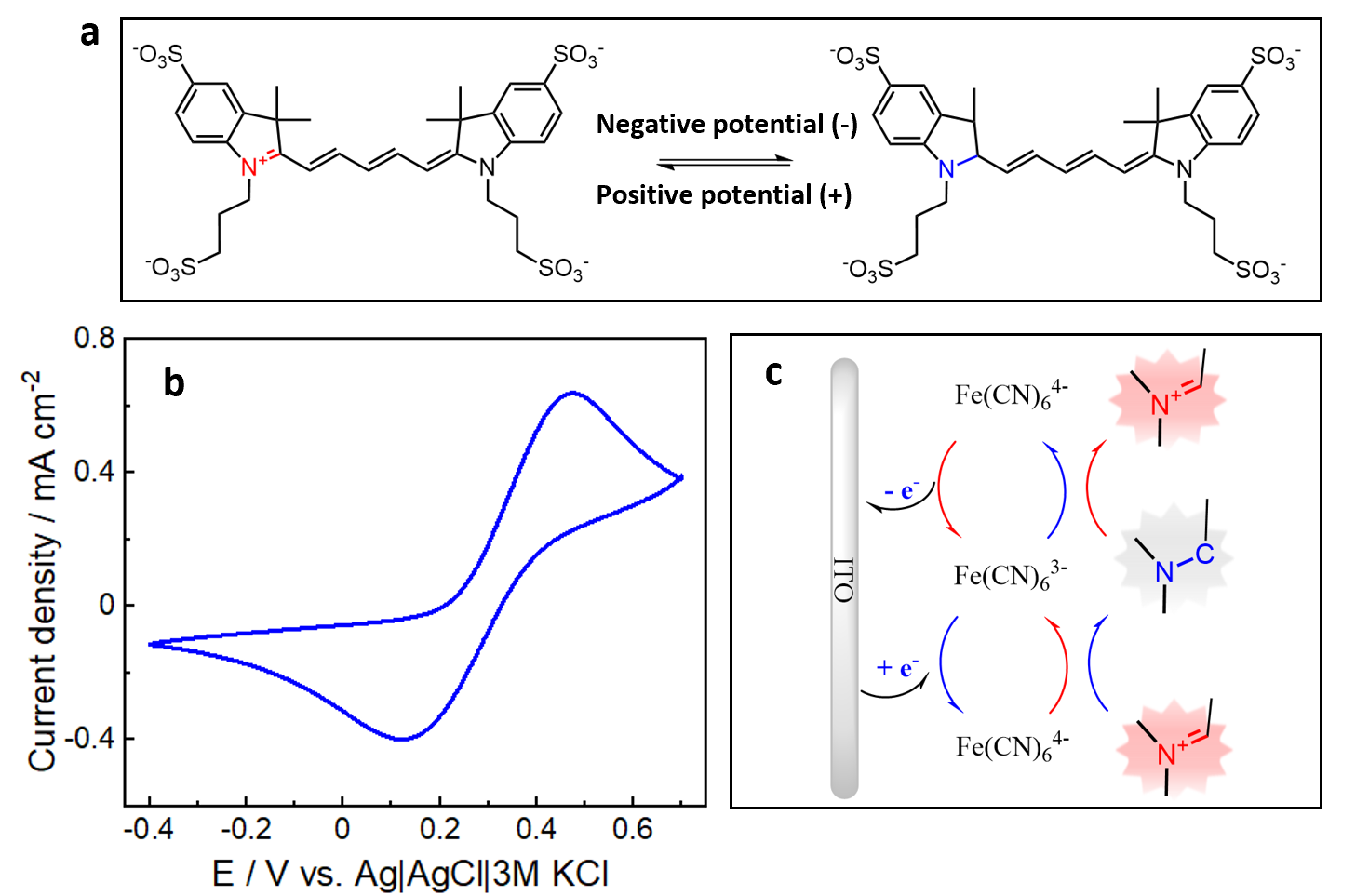

**Supplementary Figure 3. a,** Electrochemical redox reaction of Alexa 647 mediated by Trolox or ferricyanide, the redox reaction happens between the quaternary ammonium cation and a tertiary amine. **b**, Cyclic voltammetry for Alexa 647/ITO in oxygen scavenger tris imaging buffer (see Method) with 1 mM ferricyanide, scan rate is 50 mV / s. **c** Scheme representation of the sequence of the ferricyanide mediated Alexa 647 redox reactions during cyclic voltammetry.

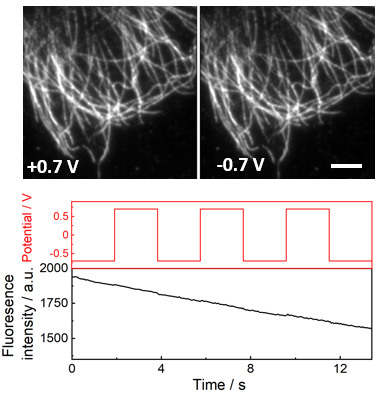

**Supplementary Figure 4.** Alexa 647 shows no obvious fluorescence intensity change by switching the applied potential at ITO surface. The imaging buffer is oxygen scavenger tris buffer without adding Trolox, ferricyanide, or cysteamine. A 642 nm laser with intensity of 0.1 kW cm^-2^ was on during the imaging. All the Alexa 647 stays in the bright state when the electrochemical potential was cycled between 0.7 V and -0.7 V. The decay in fluorescence intensity was due to photobleaching during the images acquisition. The scale bar is 5 μm.

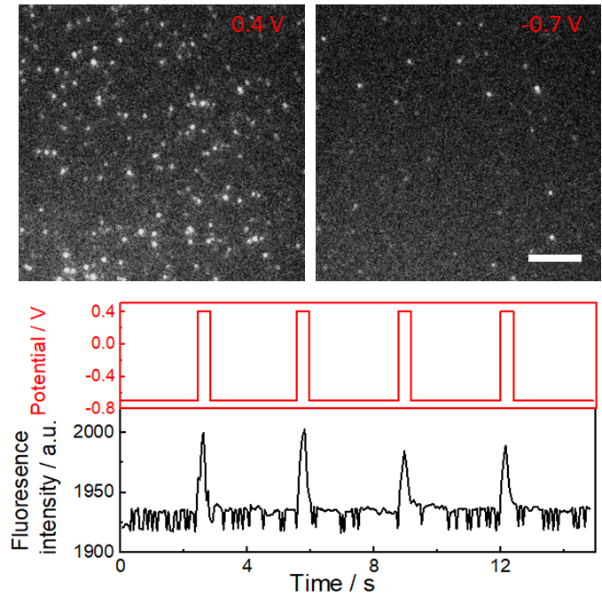

**Supplementary Figure 5.** TIRF images of Alexa 647 molecule adsorbed onto poly-L-lysine modified ITO slide in oxygen scavenger tris buffer with 50 mM cysteamine. The corresponding electrochemical potential regulated fluorescence switching curve was shown. The majority of Alexa 647 are in their ON state at positive potentials (0.4 V) and turned off to the dark state at negative potentials (-0.7 V). A 642 nm laser with intensity of 2 kW cm^-2^ was used as the excitation source during imaging**.** The scale bar is 20 μm.

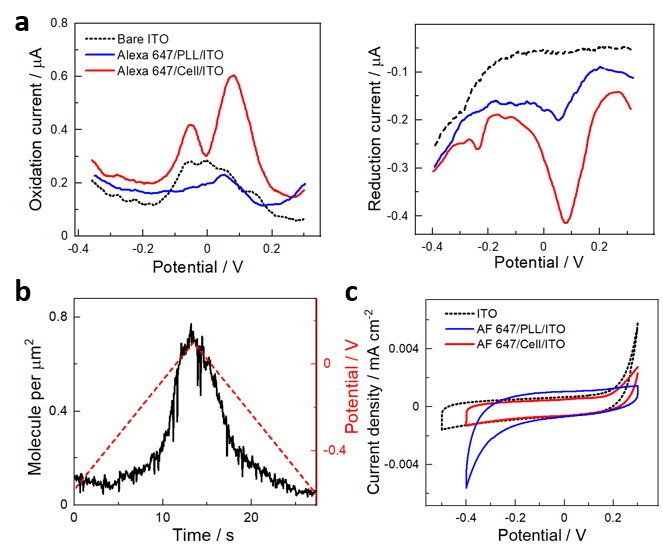

**Supplementary Figure 6. a,** Different pulsed voltammetry curves for ITO (black dotted line), Alexa 647 adsorbed onto polylysine coated ITO (Alexa 647/PLL/ITO, blue line), and Alexa 647 labelled COS-7 sample on ITO (Alexa 647/Cell/ITO, red line) in a STORM buffer. **b,** For Alexa 647 attached polylysine coated ITO (Alexa 647/PLL/ITO) sample, The ON state Alexa 647 increases with more positive applied potentials during cyclic voltammetry scan in STORM buffer. **c**, Cyclic voltammetry for ITO (black dotted line), Alexa 647 attached polylysine coated ITO (Alexa 647/PLL/ITO, blue line), and Alexa 647 labelled COS-7 sample on ITO (Alexa 647/Cell/ITO, red line) in oxygen scavenger tris imaging buffer without adding Trolox, ferricyanide, or cysteamine.

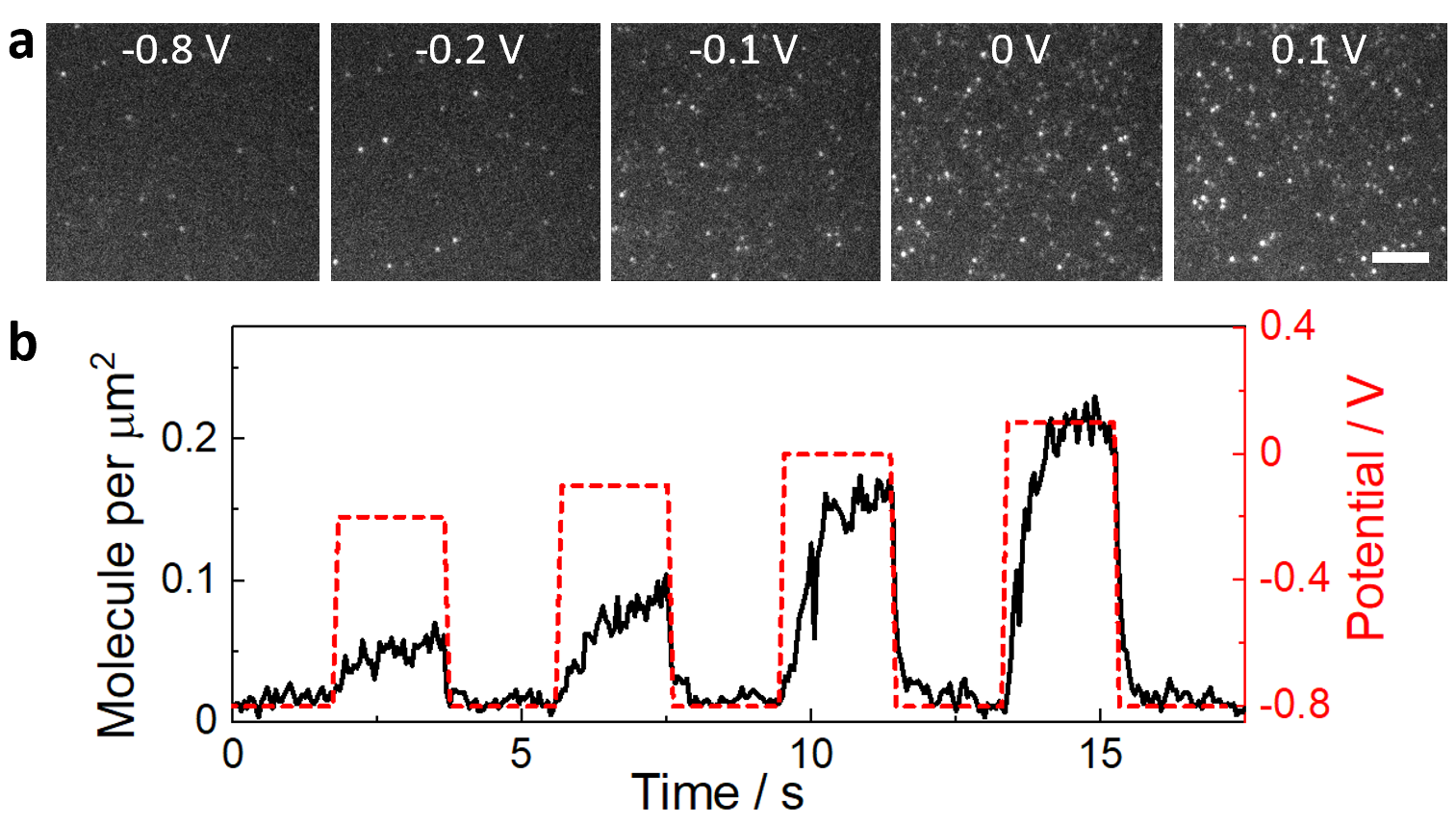

**Supplementary Figure 7. a,** TIRF images of Alexa 647 molecule adsorbed onto poly-L-lysine modified ITO slide in oxygen scavenger tris buffer with 50 mM cysteamine under various electrochemical potentials. A 642 nm laser with intensity of 2 kW cm^-2^ was used as the excitation source during imaging**.** The scale bar is 20 μm. **b**, The ON-state Alexa 647 density increases with applied potentials.

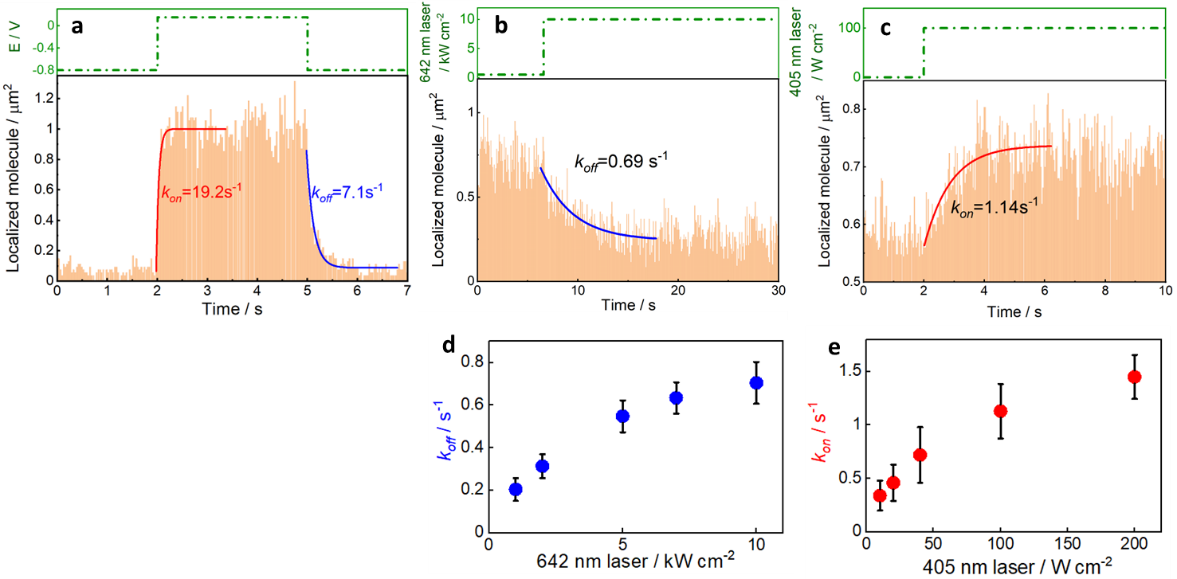

**Supplementary Figure 8. a,** Measurement for the rate constant *k_on_* and *k_off_* using the electrochemical switching method. First, the Alexa 647 labelled tubulin samples were imaged using a 642 nm reading laser (3 kW cm^-2^). By applying a negative potential (-0.8 V) to the ITO surface, Alexa 647 will be switched into the OFF state, followed a positive potential (0.2 V) will reversibly switch the Alexa 647 back to ON state. The rate constant was determined by counting the number of molecules in the field of view as function of time and then fitting the distribution to a single exponential function. For comparison, the rate constant for *k_on_* and *k_off_* using photochemical switching was measured by (**b**) irradiated the sample with strong 642 nm laser (10 kW cm^-2^) to switch Alexa 647 off (to obtain *k_off_*) first, and then reduce the 642 nm laser to 3 kW cm^-2^,(**c**) followed by turning UV light on the OFF state fluorophores are switched back to ON state (to determine *k_on_*). The rate constants *k_on_* and *k_off_* as a function of the photochemical switching conditions are summarized in **d-e**. Error bars in d and e denote mean ± standard deviation (n = 6 replicates of distinct samples).

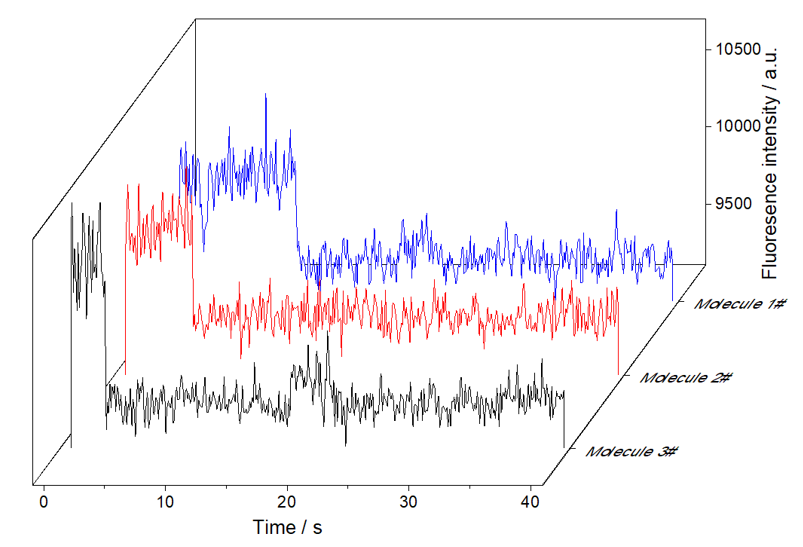

**Supplementary Figure 9.** Examples of photobleaching curves of single Alexa 647 molecules covered ITO in PBS buffer. A total of 400 frames TIRF images were taken, with exposure time of 100 ms, the 642 nm laser intensity was 1 kW cm^-2^. By analyzing the fluorescence trajectories of 411 molecules in the field of view, 100% of the molecules have only one single bleaching step. The fluorescence statistics data showed the average fluorescence background is 9491 ± 106 a.u., and the average fluorescence intensity of single Alexa 647 molecules is 10112 ± 303 a.u.

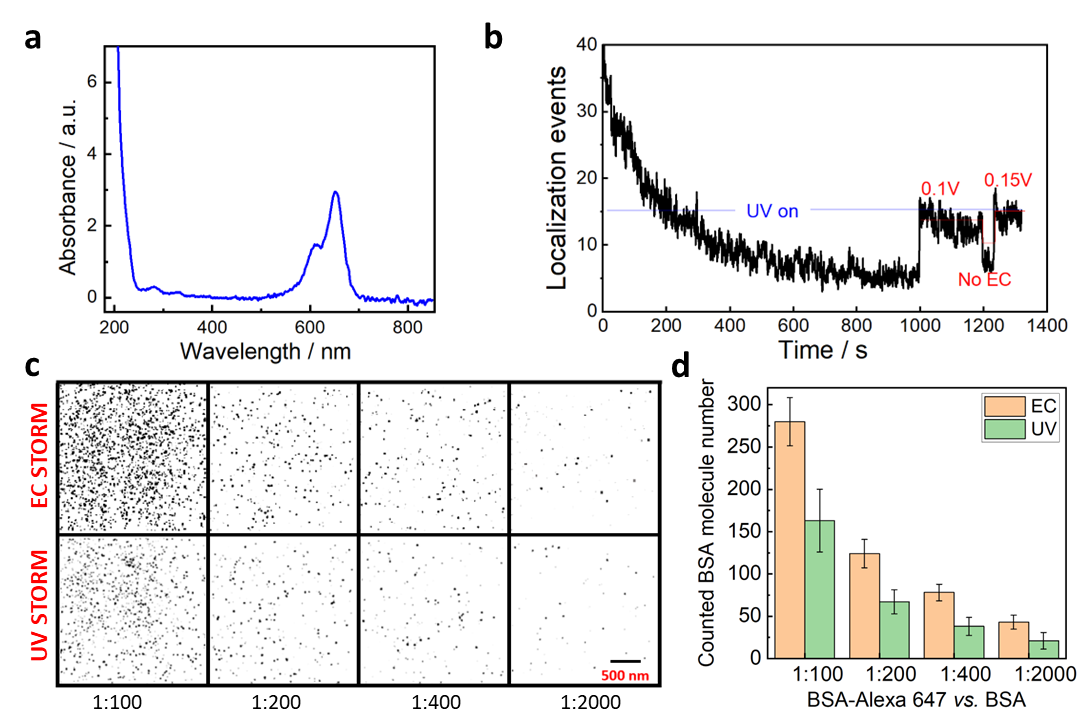

**Supplementary Figure 10.** **a**, The UV-Vis absorbance spectrum for Alexa 647 labelled BSA. The calculated degree of labelling is 2.46 (with absorbance of 2.74 at the wavelength of 280 nm and absorbance of 28.55 at the wavelength of 655 nm). **b**, A BSA-Alexa 647 sample where the conventional combination of lasers (4 kW cm^-2^ for 642 nm laser and 100 W cm^-2^ for 405 nm laser) are employed to exciting Alexa 647 for STORM imaging at the beginning of the imaging session. After image acquiring for 600 s (12,000 frames, with frame exposure time of 50 ms) there are very few emitters. A positive potential of 0.1 V was applied at 1000 s which turned many of the OFF state fluorophores back to the ON state. During 1000 to 1300 s, it can be seen that the positive electrochemical potential can effective turn more off-state Alexa 647 to on state than using UV laser activation only. **c**, Direct STORM images for BSA-Alexa 647 at ITO surface under electrochemical (top panels) and laser-controlled (bottom panels) Alexa 647 blinking mode. From left to right panels, to create gradient BSA-Alexa 647 covered surface, the BSA-Alexa 647 were diluted with BSA with ratios of 1:100, 1:200, 1:400, 1:2000, respectively, before attaching them on ITO surface. Under ‘EC’ conditions, no UV laser was used and rather positive potentials were applied to the ITO surface in a range between 0.1 to 0.35 V during data collection to ensure as many Alexa 647 could be activated to the ON state as possible. Under ‘UV’ condition, the UV laser intensity was increased from 10 to 200 W cm^-2^, and no electrochemical potential was applied to the ITO. **d**, Recovery yield comparison using BSA-Alexa 647 sample (see Method for sample preparation) under the same ‘UV’ or ‘EC’ conditions as described in **c**. Error bars in f denote mean ± standard deviation (n = 9 replicates).

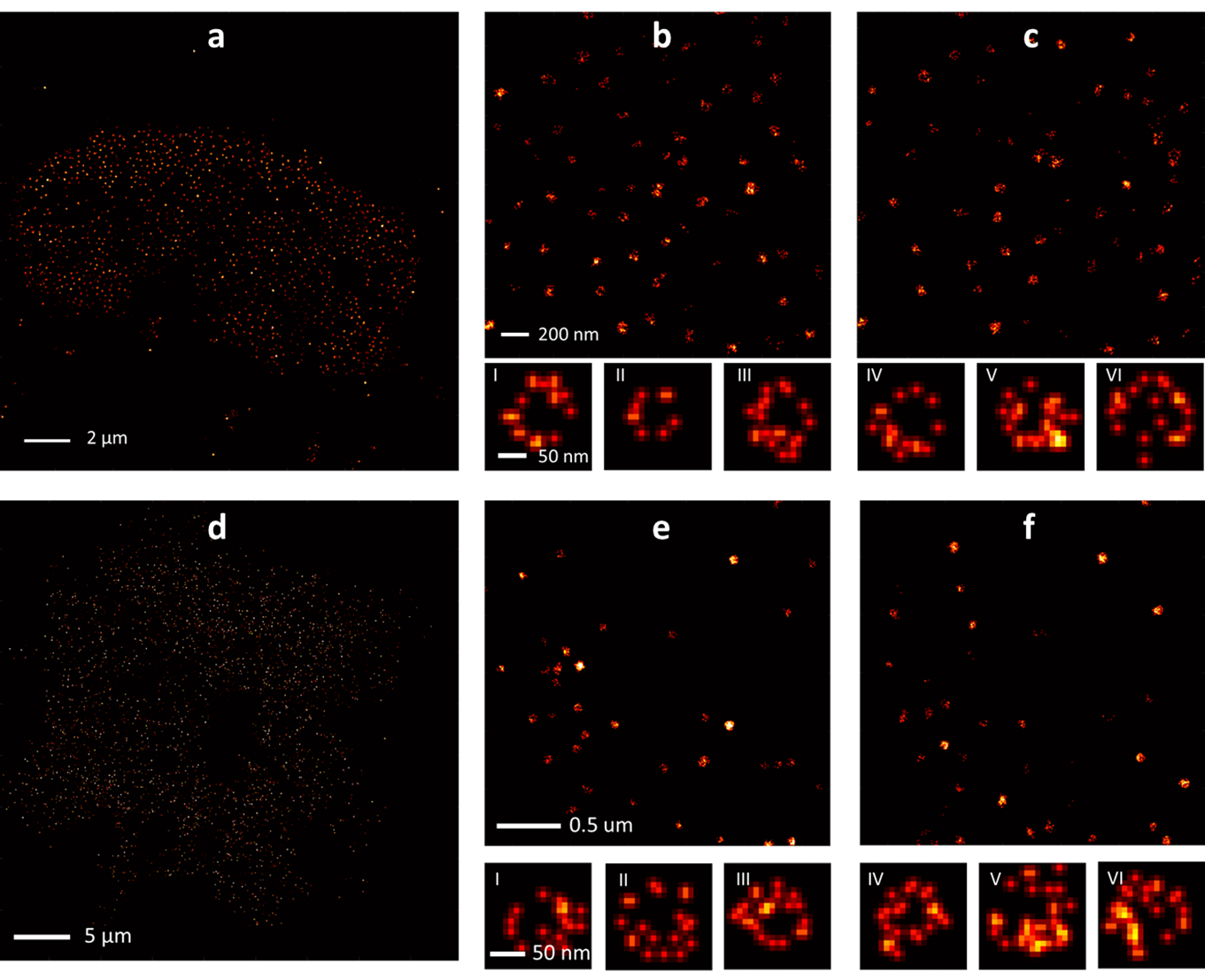

**Supplementary Figure 11. STORM images of nuclear pore complex and clathrin-coated pits.** **a,** EC-STORM of the cell nucleus of a COS-7 cell stained with primary anti-Nup98 antibodies and Alexa 647 conjugated secondary antibodies. Zoom in EC-STORM (**b**) and conventional STORM (**c**) images of the same area of one cell nucleus both revealed the ring like structures of nuclear pore complex, further magnified EC-STORM (I-III) and conventional STORM (IV-VI) images revealed the symmetrical and octagonal arrangement of Nup98 within the nuclear pore complex. **d,** EC-STORM of a large area of one COS-7 cell stained with primary anti-clathrin antibodies and Alexa 647 conjugated secondary antibodies. EC-STORM (**e**) and conventional STORM (**f**) images of same area of one COS-7 cell resolved the spherical structure of matured clathrin-coated pits with zoomed in examples (I-III for EC-STORM, IV-VI for conventional STORM). In ‘conventional STORM’ condition, 4-6 kW cm^-2^ 642 nm laser and 0-0.5 kW cm^-2^ UV lasers was altered in real time according to the emitter density change, ‘EC-STORM’ condition refers to 4 kW cm^-2^ 642 nm laser and slowly increasing the applied potential from -0.3 V to 0.1 V during image acquisition.

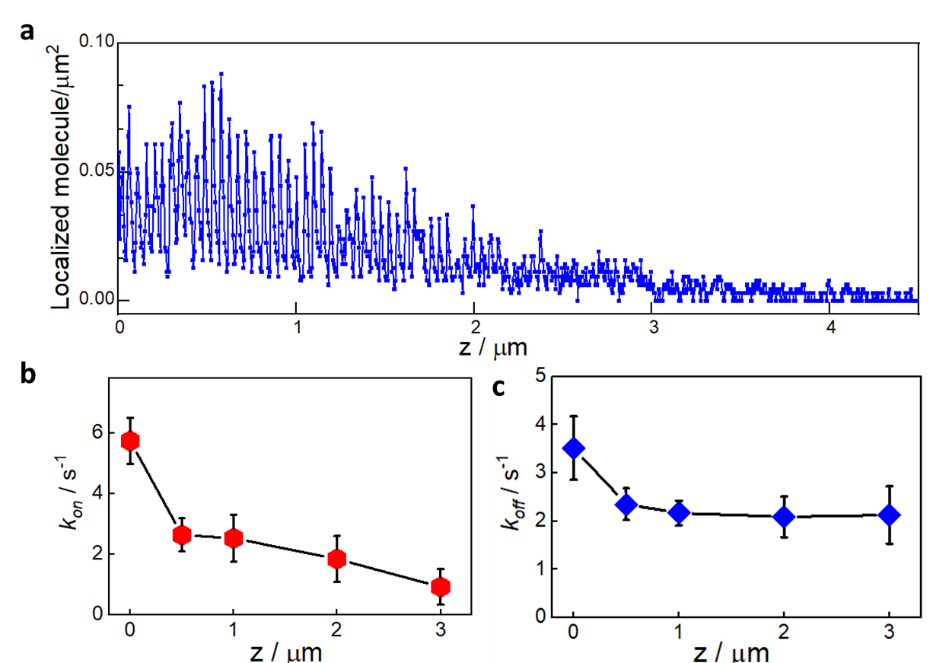

**Supplementary Figure 12.** The change for Alexa 647 switching capability using EC-STORM with the distance increase from the ITO surface. **a**, When an Alexa 647 labelled polystyrene beads (with diameter of 100 μm) are placed on an ITO electrode in STORM buffer, the Alexa 647 on the beads shows the switching response to the electrochemical potential, and this response becomes slower with the distance increase between Alexa 647 and ITO surface. The electrochemical potential was switched between 0.1 V for 0.2 s and -0.8 V for 0.5 s for multiple cycles. The corresponding movie (Supplementary Movie 7) shows the dynamic switching with the electrochemical potential switching. **b-c**, In high inclined and laminated optical sheet (HILO) illumination mode, the *k_on_* and *k_off_* using EC-STORM were obtained at a list of z distances from ITO.

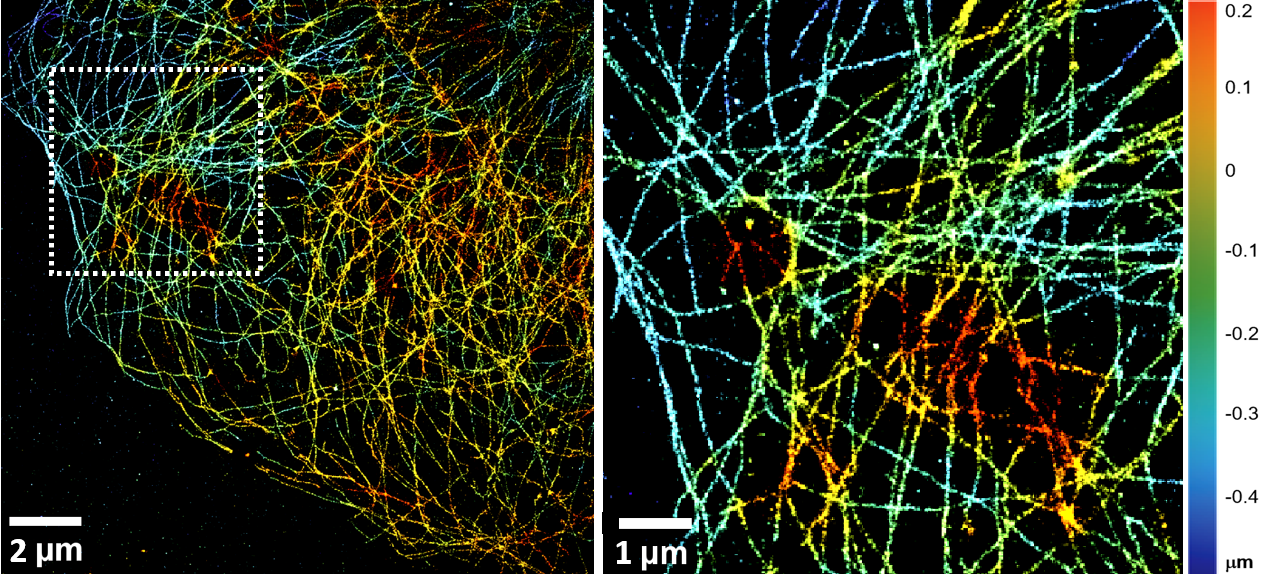

**Supplementary Figure 13.** 3D EC-STORM image of cellular microtubule network on COS-7 cells. The axial position is represented as a z-colour coded map. Image was reconstructed from 30000 frames of 33 ms exposure time to reach a localization precision ∼25 nm along the radial direction and ∼50 nm along the axial one.

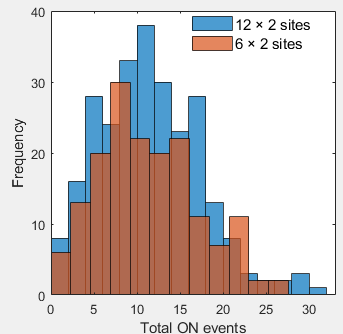

**Supplementary Figure 14**. Histogram of the total ON events of Alexa647 on two groups of nanorulers (with 6 × 2 and 12 × 2 binding sites for Alexa 647) by UV light activation. 1 kW cm^-2^ of 642 nm laser and 10 W cm^-2^ of UV laser were on during imaging.

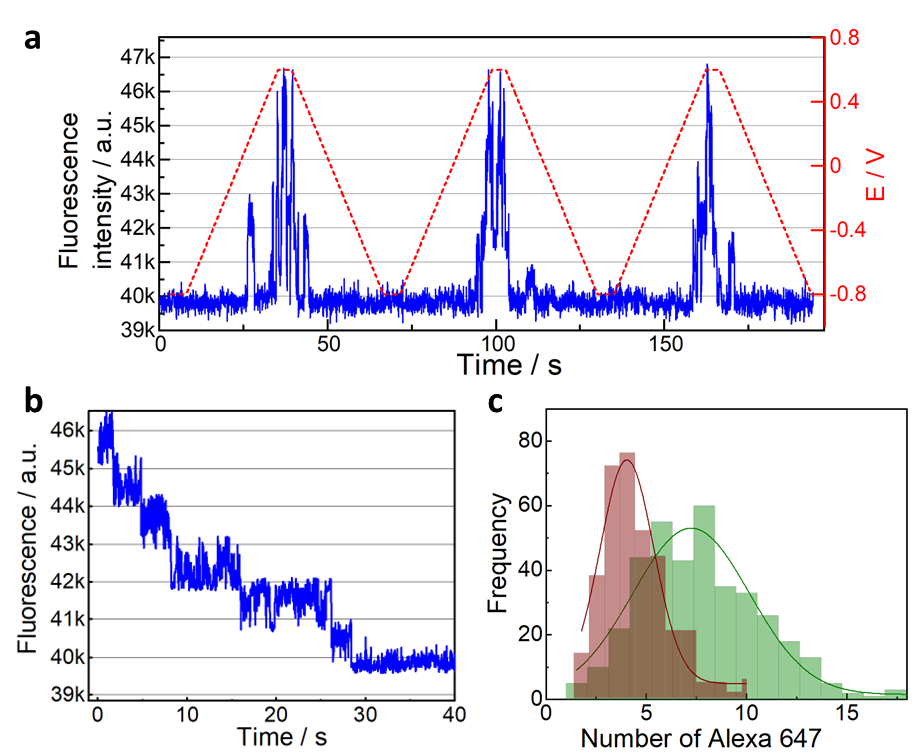

**Supplementary Figure 15**. **a**, How the Alexa 647 molecules can be reversibly turned on and off at one nanoruler (blue line for stepwise fluorescence intensity change) by linearly increasing or decreasing the electrochemical potential (red dashed line) with a scan rate of 50 mV s^-1^. The applied potential was held for 2 s at -0.8 V and 0.6 V before scanning potential progressively to more positive or negative values each time. **b**, By regrouping the segmented fluorescence traces (from panel e) with similar intensity and ranking the grouped segmented traces in descending order, a stepwise fluorescence trajectory curve was obtained. **c,** Histogram for the number of Alexa 647 counted at each DNA nanoruler by taking the ratio of the maximum intensity obtained in **a** to the single molecule brightness of Alexa647 obtained by step fitting in **b**.

**Supplementary Movie 1.** The electrochemical potential modulated fluorescence switching of Alexa 647 in oxygen scavenger tris buffer with 1 mM ferricyanide. The imaging sample is COS-7 cells labeled for microtubules with Alexa 647.

**Supplementary Movie 2**. The electrochemical potential modulated fluorescence switching of Alexa 647 in oxygen scavenger tris buffer with 50 mM cysteamine, while at constant potential, Alexa 647 shows stochastic blinking property. The imaging sample is COS-7 cells labeled for actin with Alexa 647.

**Supplementary Movie 3.** The ON state Alexa 647 molecule density changes during cyclic voltammetry scan. During the imaging session, the electrochemical potential was ramped from -0.4 V to 0.3 V and then back to -0.4 V, with scan rate of 20 mV s^-1^. The imaging sample is COS-7 cells labeled for microtubules with Alexa 647, and imaging buffer is oxygen scavenger tris buffer with 50 mM cysteamine.

**Supplementary Movie 4.** The ON state Alexa 647 molecule density when the electrochemical potentials are switched between several relative positive values (-0.4, -0.2, -0.1, 0, and 0.1 V) and -0.7 V for 10 s at each still potential. The imaging sample is COS-7 cells labeled for microtubules with Alexa 647, and imaging buffer is oxygen scavenger tris buffer with 50 mM cysteamine.

**Supplementary Movie 5.** The ON state Alexa 647 molecule density at different positive potentials as indicated. The imaging sample is COS-7 cells labeled for microtubules with Alexa 647, and imaging buffer is oxygen scavenger tris buffer with 50 mM cysteamine.

**Supplementary Movie 6.** The ON state Alexa 647 molecule density was increased dramatically by increasing the power of 642 nm laser, while a negative potential could dramatically reduce the emitter density. After releasing the negative potential, under open circuit potential, the ON state Alexa 647 molecule slowly increased.

**Supplementary Movie 7.** When an Alexa 647 labelled polystyrene beads (with diameter of 100 μm) are placed on ITO electrode in STORM buffer, the Alexa 647 on the beads shows the switching response to the electrochemical potential. The electrochemical potential was switched between 0.1 V for 0.2 s and -0.8 V for 0.5 s for multiple cycles. The movie was recorded under high inclined and laminated optical sheet (HILO) illumination mode, with frame rate of 50 ms.

**Supplementary Movie 8.** The ON and OFF switching of Alexa 647 at DNA nanoruler sample (two ends of the DNA nanoruler has 6 binding sites for Alexa 647 and that were separated by 94 nm) under EC-STORM where the potential was pulsed. 1 kW cm^-2^ of 642 nm laser was on, the electrochemical potential was pulsed between -0.6 V for 900 ms and 0.1 V for 100 ms for multiple cycles.

References

1.	J. S. Mayell, A. J. Bard, The electroreduction of quaternary ammonium compounds. *J. Am. Chem. Soc.* **85**, 421-425 (1963).

2.	S. Fan *et al.*, Observing the Reversible Single Molecule Electrochemistry of Alexa Fluor 647 Dyes by Total Internal Reflection Fluorescence Microscopy. *Angew. Chem., Int. Ed.* **131**, 14637-14640 (2019).

3.	M. G. Gustafsson, Surpassing the lateral resolution limit by a factor of two using structured illumination microscopy. *J. Microsc.* **198**, 82-87 (2000).

4.	M. G. Gustafsson, Nonlinear structured-illumination microscopy: wide-field fluorescence imaging with theoretically unlimited resolution. *Proc. Natl. Acad. Sci.* **102**, 13081-13086 (2005).

5.	M. F. Juette *et al.*, Three-dimensional sub–100 nm resolution fluorescence microscopy of thick samples *Nat. Methods* **5**, 527-529 (2008).
